# Intratumoral Treg ablation is sufficient to mediate tumor control systemically without autoimmunity

**DOI:** 10.1101/2025.06.26.661765

**Authors:** Alissa Bockman, Brenna Gittins, Chenyu Zhang, Jessica Hung, Mariela Moreno Ayala, Dorothée Saddier Axe, Brian M Weist, Michel DuPage

## Abstract

Regulatory T cells (T_regs_) infiltrate most tumors, yet whether they suppress immune responses directly within tumor tissues is not clear. We used intratumoral (IT) delivery of diphtheria toxin (DT) in *Foxp3^DTR^* mice to deplete IT T_regs_ while leaving peripheral T_regs_ intact. IT delivery of DT reduced T_reg_ frequencies in the tumor, which promoted potent tumor control without autoimmunity. Interestingly, this control was principally mediated by CD4^+^ T cells, whereas CD8^+^ T cells only contributed when CD4^+^ T cells were absent. While conventional dendritic cells (cDCs) were required to clear tumors, Batf3^+^ cDC1s were dispensable. Distant secondary tumors, mimicking metastases, were also controlled by IT T_reg_ ablation. Mechanistically, IT T_regs_ suppressed antitumor T cell responses by blocking the acquisition of tumor antigen by cDC2s. Importantly, similar mechanisms of control were observed using a clinically translatable IT T_reg_-depleting anti-CCR8 antibody and reveal a distinct therapeutic strategy that leverages cDC2s and CD4^+^ T cells.

## INTRODUCTION

High frequencies of tumor-infiltrating regulatory T cells (T_regs_) are correlated with reduced survival in cancer patients^1–6^. T_regs_ inhibit immune responses through the secretion of suppressive cytokines and the inhibition of costimulatory ligand expression on APCs, among other suppressive mechanisms^7–10^. Eliminating T_regs_ as a therapy is complicated by the fact that T_regs_ are essential for preventing harmful inflammation, including autoimmunity^11–14^. Therefore, to target T_regs_ for the treatment of cancer will require approaches that only disrupt T_reg_ suppression of antitumor immunity while retaining T_reg_ immunoregulatory function in non-tumor tissues.

One approach would be the selective depletion of T_regs_ within tumors, but studies have suggested that T_regs_ may suppress antitumor immune responses outside of the tumor microenvironment^15–19^. Nevertheless, antibody-based strategies that selectively deplete intratumoral (IT) T_regs_ have worked well in mouse models, for example, with antibodies that can deplete intratumoral T_regs_ by targeting CTLA-4, CD25, GITR, 4-1BB, OX-40, LAG3, TIGIT orCCR4^5,10,20–275,10,20–26^. Most recently, T_regs_ were found to upregulate the chemokine receptor CCR8 in tumors from humans and mice^28–32^, leading to experiments showing that anti-CCR8 antibodies can effectively control several types of cancer in mice^33–38^. Currently, phase I and II clinical trials with anti-CCR8 are underway for advanced and metastatic solid cancers (NCT05537740; NCT05101070; NCT06387628; NCT05007782; NCT05635643; NCT05935098). Despite the promise of these new antibody approaches that can selectively deplete IT T_regs_, the fundamental question as to whether T_regs_ suppressive function occurs only in tumors is not resolved. That is because such antibody approaches cannot be solely restricted to tumor tissues, as cells outside of tumors expressing the target antigen (i.e. CCR8), which include non-T_reg_ cells, may also contribute to tumor control^39–41^. Furthermore, the activation of Fc receptors on immune cells that engage therapeutic antibodies may be critical for the efficacy of these therapeutics^27,42–4442–44^. Therefore, further research into T_reg_ depletion without using antibodies will directly test whether IT T_reg_ ablation alone directs anti-tumor immunity.

To prevent the activity of antibodies binding to cells outside of the tumor microenvironment, investigators have performed IT delivery of antibodies, as well as immune adjuvants CpG and IL-12-Fc^45–48^. Direct, intratumoral TLR9 activation by CpG injection promoted the control of treated tumors as well as untreated tumors^45^. When TLR9/CpG administration was combined with anti-CTLA4 or anti-OX40 to deplete IT T_regs_, therapeutic activity was enhanced, leading to effective control of both treated tumors as well as disseminated tumors^45–48^. Intratumoral immunotherapy poses significant clinical benefit, as local delivery may mitigate the autoimmune toxicities that are elicited by systemic drug administrations^49^. However, the role of Fc receptor binding and immune cell activation are still likely to be important, and alternative methods that do not employ antibodies to remove intratumoral T_regs_ must be tested^50^.

To directly test whether T_reg_ ablation only within tumors is sufficient to control tumors without inciting autoimmunity, we delivered a low dose of diphtheria toxin directly into tumor tissues by intratumoral diphtheria toxin injection (IT DT) in *FoxP3^DTR^*mice^11,51^. This technique led to nearly complete ablation of IT T_regs_ while sparing T_regs_ at peripheral sites throughout the mice. While others have used higher doses of IT DT to ablate tdLN and IT T_regs_, here we establish a lower dose DT strategy in which IT T_regs_ are ablated with minimal tdLN T_reg_ depletion^27^. Most importantly, IT T_reg_ ablation by itself led to potent tumor control without inciting autoimmune disease. Furthermore, the ablation of T_regs_ from a single tumor led to control of distant tumors of the same origin that did not receive direct treatments. Unexpectedly, tumor control was dominantly mediated by CD4^+^ T conventional (Tconv) cells rather than CD8^+^ T cells, although CD8^+^ T cells did contribute to tumor control in the absence of CD4^+^ T cells. In addition, cDC1s were not required for tumor control with IT T_reg_ ablation. Mechanistically, we show that IT T_regs_ suppressed the antitumor response locally by blocking the acquisition of tumor antigen by cDC2s within tumors. We demonstrate enhanced antigen acquisition by cDC2s upon the removal of IT T_reg_ suppression. Similar cell type dependencies were observed with clinically applicable anti-CCR8 intratumoral T_reg_ depleting antibodies. These findings are consequential, as they uncover a clinically relevant mechanism to boost systemic antitumor immune responses that does not rely on CD8^+^ T cells or cDC1s and is effective without inciting autoimmune toxicity.

## RESULTS

### Intratumoral T_reg_ depletion controls tumor growth without inciting autoimmunity

It is unclear whether approaches to deplete intratumoral (IT) T_regs_ selectively would have clinical efficacy, as systemic T_reg_ depletion has almost exclusively been tested^52–54^. Several reports have shown that T_reg_ depletion outside of the tumor, e.g. in tumor-draining lymph nodes, is critically important for tumor control^15,16,18,55^. However, recent work has also shown that anti-CCR8 antibodies, which deplete T_regs_ selectively within tumors, prolong mouse survival after challenge with subcutaneously implanted CT26 colorectal carcinoma or EMT6 breast cancer cell lines^33,34^. We confirmed previous findings that T_regs_ within MC38 tumors highly express CCR8, and that anti-CCR8 treatment reduces CCR8^+^ IT T_reg_ numbers in tumors and significantly decreases MC38 tumor burden (Fig. S1A-G)^28,29,35,37^. Interestingly, anti-CCR8 treatments were not effective against B16F10 tumors (Fig. S1F-G). Overall, these data were consistent with an important role for the local activity of IT T_regs_ in mediating tumor immunotolerance. However, because CCR8 can be expressed on a subset of circulating T_regs_ and CD4^+^ T conv cells, it remained possible that anti-CCR8 therapy exerted tumor control by acting more broadly than on IT T_regs_ alone. Furthermore, antibody-mediated cell depletion that activates Fc receptors may have also contributed to the antitumor efficacy^43^. To test whether IT T_reg_ ablation alone is sufficient to promote tumor control without the use of antibodies, we used *FoxP3^DTR^* mice. However, rather than deliver diphtheria toxin (DT) systemically by intraperitoneal (IP) injection, we delivered low dose DT (250 ng, one-fourth of the dose used IP) directly into tumors (IT) every other day once tumors reached a volume of 50 mm^3^ and as long as tumors were palpable (Fig. 1A). After a single dose of IT DT, IT T_regs_ were efficiently depleted by 48 hours (Fig. 1B). Importantly, we found that IT DT ablation of IT T_regs_ induced significant control of both immune-rich MC38 tumors and poorly immunogenic B16F10 tumors. As IT DT treatment was stopped after complete tumor regression occurred, we found that no tumors rebounded upon cessation of treatment (Fig. 1C,1D).

**Figure 1.**
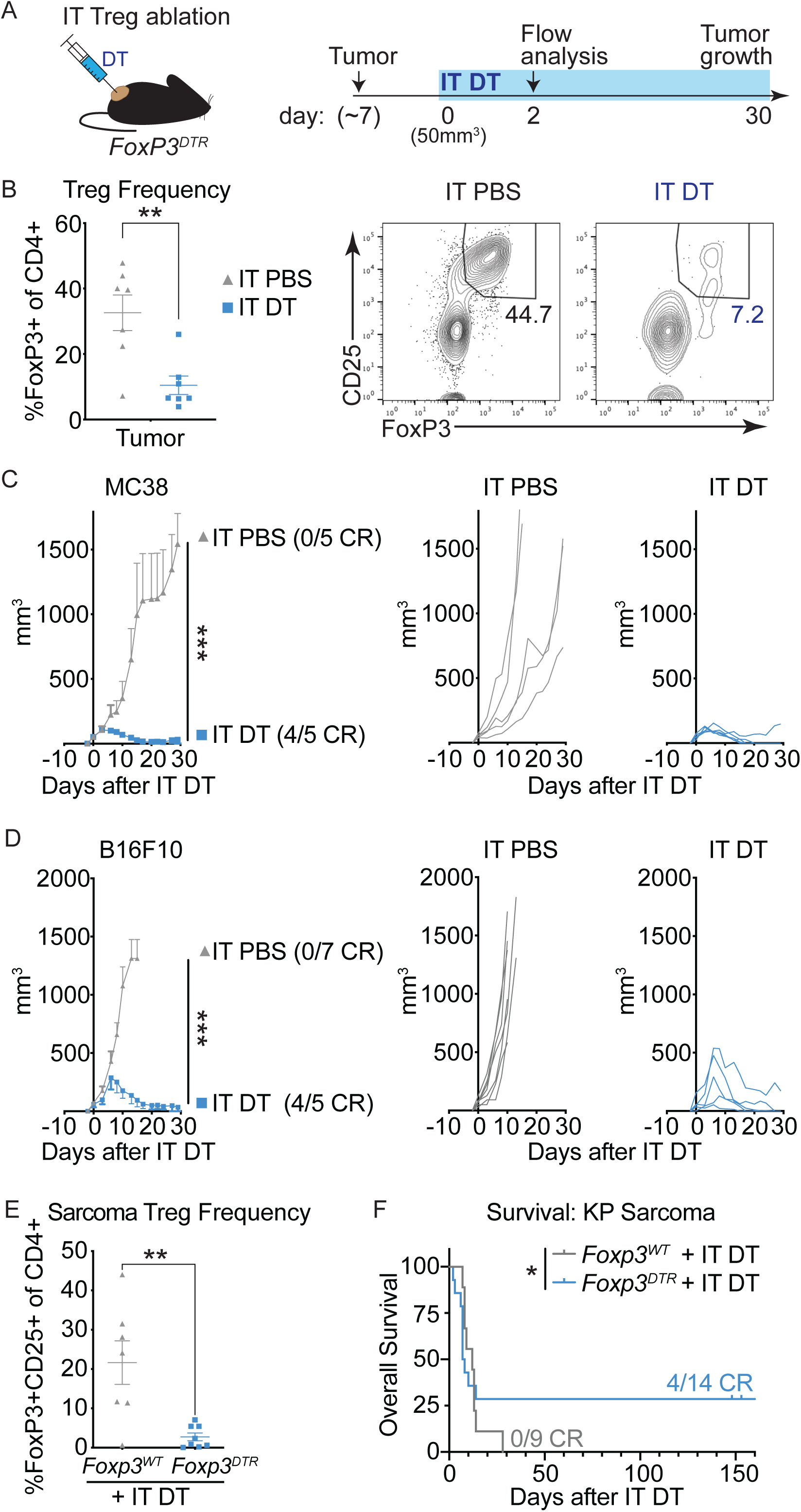
Intratumoral Treg depletion is sufficient to control tumor growth. (A) Experimental design for intratumoral diphtheria toxin (IT DT) injection in *Foxp3^DTR^* mice. DT was administered every 48-72 hours beginning when tumors were 50mm^3^ and throughout the study. (B) IT Treg frequencies in MC38 tumor-bearing *Foxp3^DTR^* mice 48 hours after one dose of IT PBS or IT DT. Representative flow cytometric staining for FoxP3+ CD25+ Tregs of live CD4+ T cells. (C-D) MC38 tumor growth (C) or B16F10 tumor growth (D) in *Foxp3^DTR^* mice after sustained IT DT or IT PBS treatment. Data are representative of 4-10 independent experiments for each tumor type. (E) IT Treg frequencies in IT DT treated *K-ras^G12D^;p53^-/-^* versus *Foxp3^DTR-GFP^;K-ras^G12D^;p53^-/-^* autochthonous sarcomas. (F) Survival analysis of sarcoma-bearing mice after IT DT treatment beginning when palpable mass was detected. Data represent means ± SEM; *p < 0.05, **p < 0.01 and ***p < 0.001. B,E: Unpaired two-tailed T test. C,D: Ordinary two-way ANOVA with Tukey’s multiple comparison test. F: One-sided Chi-square test. (n=5-14 mice/group).

To determine if IT T_reg_ ablation also promotes tumor control in autochthonous models of cancer, we induced sarcomas in *K-ras^LSL-G12D^;p53^fl/fl^* (‘KP’) mice that were bred to the *FoxP3^DTR^* allele^56^. We dosed *FoxP3^DTR^;K-ras^G12D^;p53^fl/fl^*with 50,000 infectious units of a lentivirus encoding Cre recombinase (Lenti-x) into the left calf muscle^57,58^. Over the following 150 days, we monitored mice for spontaneous tumor development. When tumors became palpable, mice were administered IT DT as with subcutaneous tumors. We found that IT DT resulted in a significant reduction of IT T_regs_ in this autochthonous tumor setting (Fig. 1E). Ultimately, this reduction in IT T_regs_ was sufficient to extend animal survival, despite the aggressive nature of this tumor model (Fig. 1F). Importantly, Lenti-x-induced sarcomas lack neoantigens recognized by CD8^+^ T cells and were previously categorized as poorly immunogenic^57,58^. Therefore, IT T_reg_ ablation appeared to promote immune control in a manner independent of a CD8^+^ T cell response. These results demonstrate a tumor-protective function of IT T_regs_ that is conserved across multiple cancer models.

While we observed robust control of IT T_reg_ ablated tumors, we sought to address whether IT T_reg_ ablation also prevented the induction of autoimmune pathologies observed with systemic T_reg_ ablation^11–14,59^. We directly compared *FoxP3^DTR^* tumor-bearing mice treated with IT DT versus IP DT for a period of 28 days by measuring T_reg_ frequencies, autoimmune pathologies, and the activation status of conventional T cells. First, IP DT treatment induced a significant reduction in T_reg_ frequencies across all tissues analyzed, whereas IT DT impacted T_reg_ frequencies predominantly within the tumor tissue, albeit a moderate T_reg_ reduction was observed in the tumor-draining lymph node (tdLN) (Fig. 2A and Fig. S2A). Second, IP DT treatments significantly increased spleen weight and caused body weight loss in mice with tumors, but IT DT did not (Fig. 2B-C; Fig. S2B-C). Furthermore, analysis of hematoxylin and eosin stained sections from the lung, liver, kidney, and heart of IP DT treated mice revealed significant lymphocytic infiltration that was not observed in mice receiving IT DT (Fig. 2D-E, Fig. S2D-E). Third, flow cytometric analysis of CD4^+^ Tconv and CD8^+^ T cells from IP DT treated mice, but not IT DT treated mice, revealed robust increases in the expression of CD44, ICOS, CD25, PD1 and TIGIT in the spleen, non-draining lymph nodes (ndLN) and tdLN (Figures 2F-G; Fig. S2F-G). Interestingly, while IT T_reg_ ablation did not induce any sign of systemic autoimmunity, IT T_reg_ ablation did produce more durable and complete rejections (CRs) of tumors compared to systemic T_reg_ ablation by IP DT (Fig. S2H-I). These data demonstrate that IT T_reg_ ablation can control tumors without inducing aberrant immune activation or multi-organ autoimmune disease.

**Figure 2.**
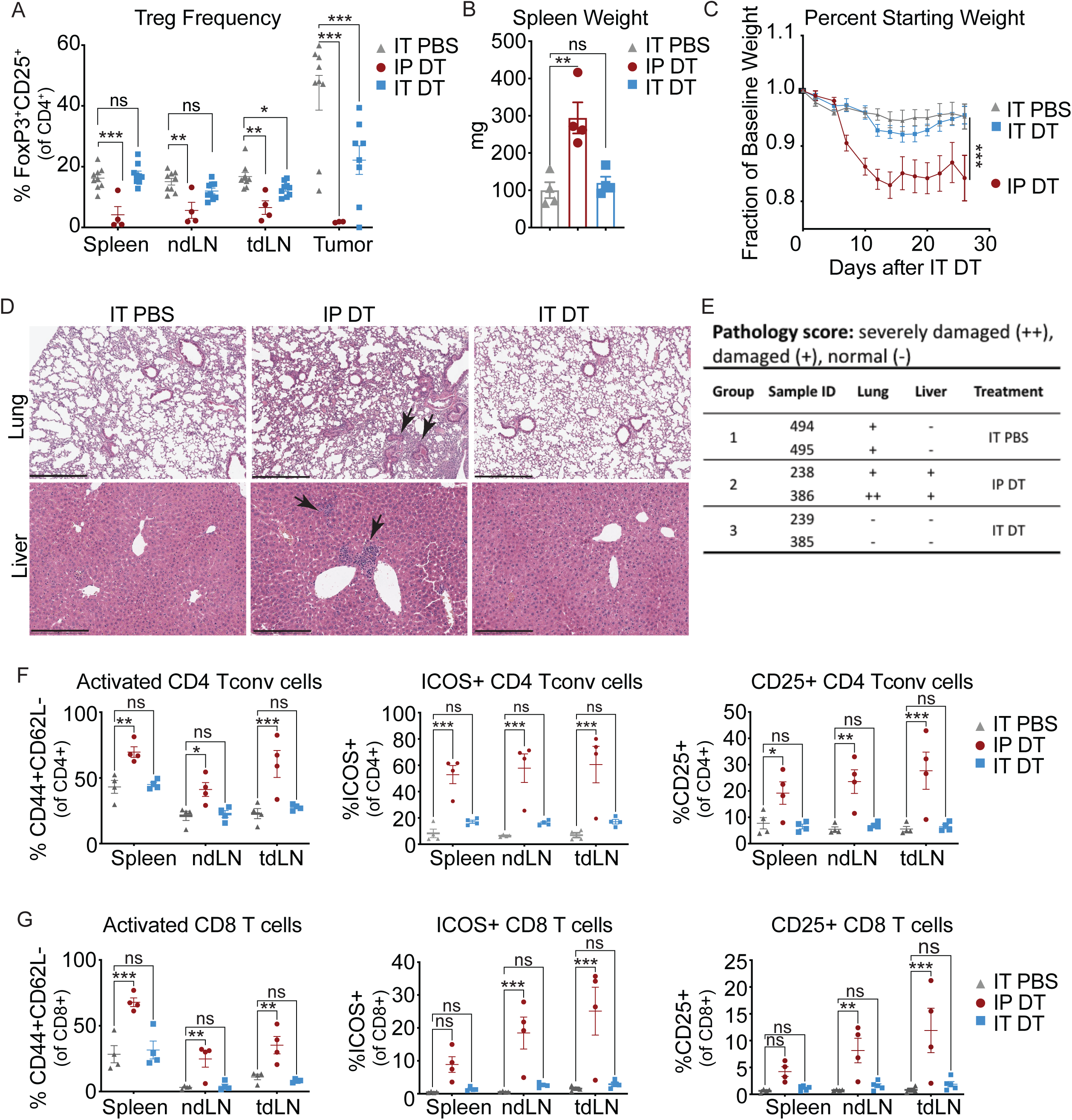
IT Treg ablation, unlike systemic Treg ablation, does not cause autoimmune pathology. (A) Treg frequencies in spleen, non-tumor-draining lymph nodes (ndLN), tumor-draining LN (tdLN), and tumor after IP PBS, IP DT, or IT DT in MC38-tumor bearing *Foxp3^DTR^* mice. (B) Spleen weight in MC38-tumor bearing *Foxp3^DTR^* mice after 30 days of sustained IP or IT DT treatment. (C) Body weight of MC38-tumor bearing *Foxp3^DTR^* mice treated with IP PBS, IP DT, or IT DT. (D) Hematoxylin and eosin staining performed on lung and liver tissues harvested after sustained IP PBS, IP DT, or IT DT. Arrows indicate lymphocytic infiltrates within tissues. Images are representative of two biological replicates for each treatment group. (E) Pathology scores of IT PBS, IP DT or IT DT treated mice. (F-G) Percent of CD4+ (F) and CD8+ (G) T cells expressing CD44, ICOS, and CD25 activation markers after sustained IP PBS, IP DT, or IT DT. Data are representative of 3 independent experiments. Data represent means ± SEM; *p < 0.05, **p < 0.01 and ***p < 0.001. A,B: Unpaired two-tailed T test. C;F;G: Ordinary two-way ANOVA with Tukey’s multiple comparison test. (n=4-9 mice/group).

### Intratumoral T_reg_ depletion promotes CD8^+^ and CD4^+^ T cell activation locally in tumors

We next characterized the immunological changes within MC38 tumors after IT T_reg_ ablation. While the frequency of CD4^+^ Tconv cells trended slightly higher with IT T_reg_ ablation, the difference was not statistically significant after 25 days of IT T_reg_ ablation (Fig. 3A). However, we did observe a significant increase in the frequency of CD8^+^ T cells, as observed by others with anti-CCR8 treatment^28,33^. To determine whether IT T_reg_ ablation also increased the frequency of tumor-reactive CD8^+^ T cells, we stained single cell suspensions collected from IT DT treated tumors with a tetramer specific for the p15E peptide of an endogenous ecotropic murine leukemia virus to detect MC38-specific CD8^+^ T cells. As has been reported for anti-CCR8 treatments^15,28,34,60–62^, we also observed a trend of an increased frequency of p15E^+^CD8^+^ T cells in tdLNs after IT T_reg_ ablation, but the increase in tumor tissues did not reach significance (Fig. S3A-B). We also noted an increase in effector (CD44+CD62L-) CD4^+^ and CD8^+^ T cells in tumor tissue after IT DT (Fig. 3B). We found that IT T_reg_ ablation induced significant upregulation of ICOS and CD25 on IT CD4^+^ Tconv cells (Fig. 3C and Fig. S3C) and CD8^+^ T cells (Fig. 3D and Fig. S3D). While the production of TNFα and IFNγ by tumor-infiltrating CD4^+^ and CD8^+^ T cells was similar between IT PBS and IT DT treated mice (Fig. 3F and Fig. S3E-I), we observed a significant increase in the frequency of IL-2 producing CD4^+^ Tconv cells after IT T_reg_ ablation (Fig. 3E and Fig. S3J). These results demonstrate that the removal of IT T_regs_ led to an increase in the activation status of conventional CD4^+^ and CD8^+^ T cells within tumor tissues.

**Figure 3.**
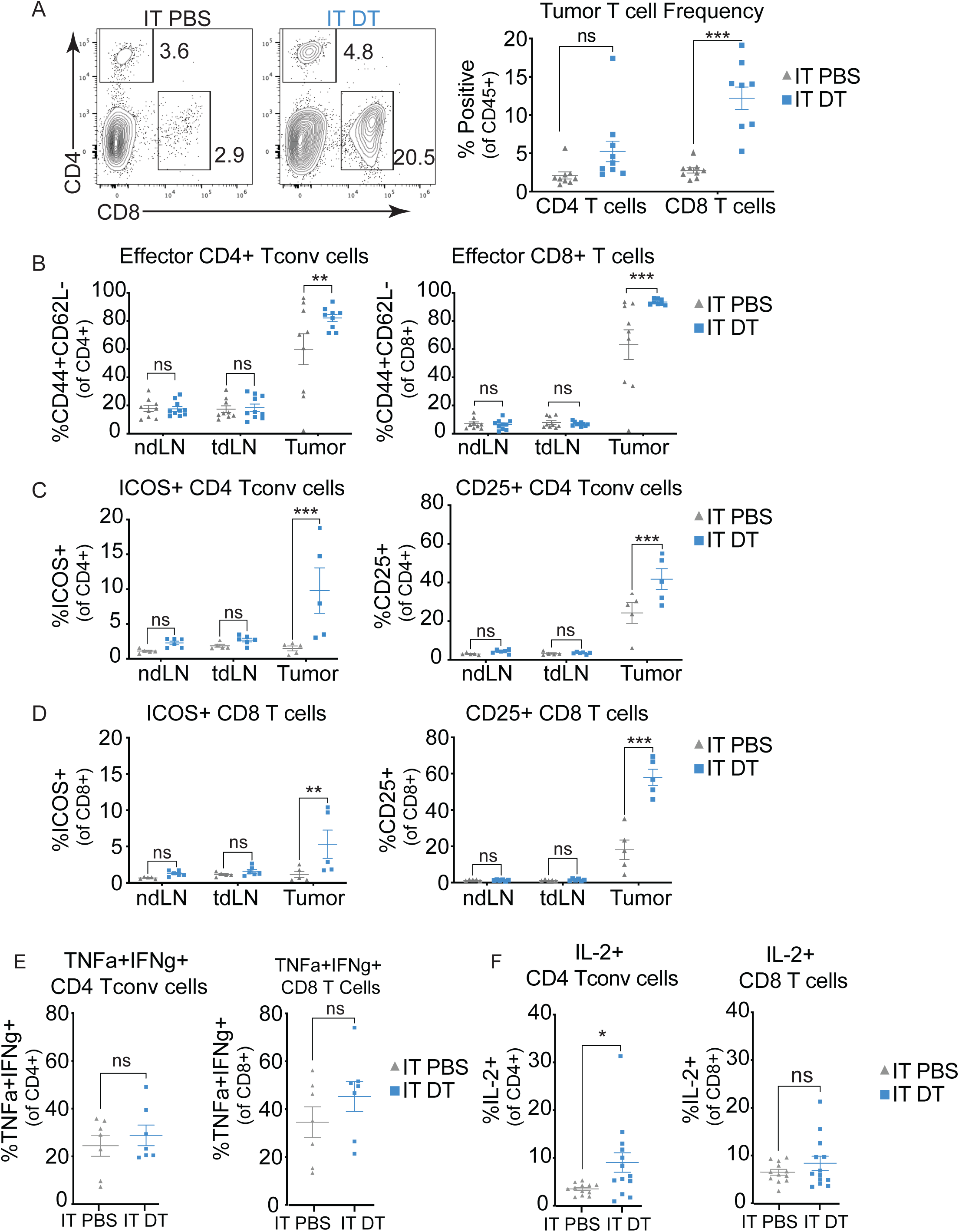
IT Treg depletion promotes CD8 and CD4 T cell activation locally in tumors. (A) Representative flow plots and quantification of CD4+ Tconv and CD8+ T cell frequencies in MC38 tumor tissue after sustained PBS or IT DT treatment. (B) Frequencies of effector (CD44+CD62L-) CD4+ Tconv cell or CD8+ T cells after IT PBS or IT DT. (C-D)Percentages of CD44+, ICOS+ or CD25+ on CD4+ Tconv cells or (D) CD8+ T cells. (E) TNF𝛼 and IFN𝛾 cytokine expression in tumor-infiltrating CD8+ and CD4+ Tconv cells. (F) IL-2 cytokine expression in CD4+ Tconv or CD8 T cells after PBS or IT DT treatment. Data are representative of 2 independent experiments. Data represent means ± SEM; *p < 0.05, **p < 0.01 and ***p < 0.001. A: Unpaired two-tailed T test. B-F: Ordinary two-way ANOVA with Tukey’s multiple comparison test. (n=5-10 mice/group).

To better understand whether the increase in T cells within tumors, observed after IT DT treatment, required the recruitment of T cells from the tdLN, we treated mice with a lymphocyte egress inhibitor (FTY720) when IT DT treatment began (D0). We found that mice treated with IT DT and FTY720 rejected tumors as efficiently as those treated with DMSO and IT DT (Fig. S3K). These data suggest that the increase in T cell frequency within IT DT treated tumors was due to the expansion and retention of T cells within tumors and not due to an influx of T cells from the tdLN. This also suggests that increased priming of T cells in lymphoid tissues is not required for tumor control with IT T_reg_ ablation. Therefore, we hypothesize that a reinvigorated T cell pool within tumor tissue itself is sufficient to promote the tumor control observed with IT T_reg_ ablation.

### CD4^+^ T conventional cells can control tumors after IT T_reg_ ablation

It is widely hypothesized that IT T_reg_ ablation unleashes CD8^+^ T cells to control tumors^15,18,21,28,33,53,60,63^. To directly test the cellular requirements for tumor control after IT T_reg_ ablation with IT DT, we performed *in vivo* cellular depletion studies using antibodies against CD4^+^ and CD8^+^ T cells or natural killer (NK1.1+) cells with IT DT administration (Fig. 4A). Tumor control and survival with IT T_reg_ ablation was unaffected by either CD8^+^ T cell or NK cell depletion alone (Fig. 4B-E and Fig. S4A-B). However, depletion of CD4^+^ Tconv cells significantly impaired tumor control and survival with IT T_reg_ ablation. Interestingly, CD8^+^ T cell depletion in combination with CD4^+^ T cell depletion fully reversed tumor control with IT T_reg_ ablation. Thus, CD4^+^ T cells played a primary role in tumor control with IT T_reg_ ablation, while CD8^+^ T cells only appeared to be required for tumor control in the absence of CD4^+^ T cells.

**Figure 4.**
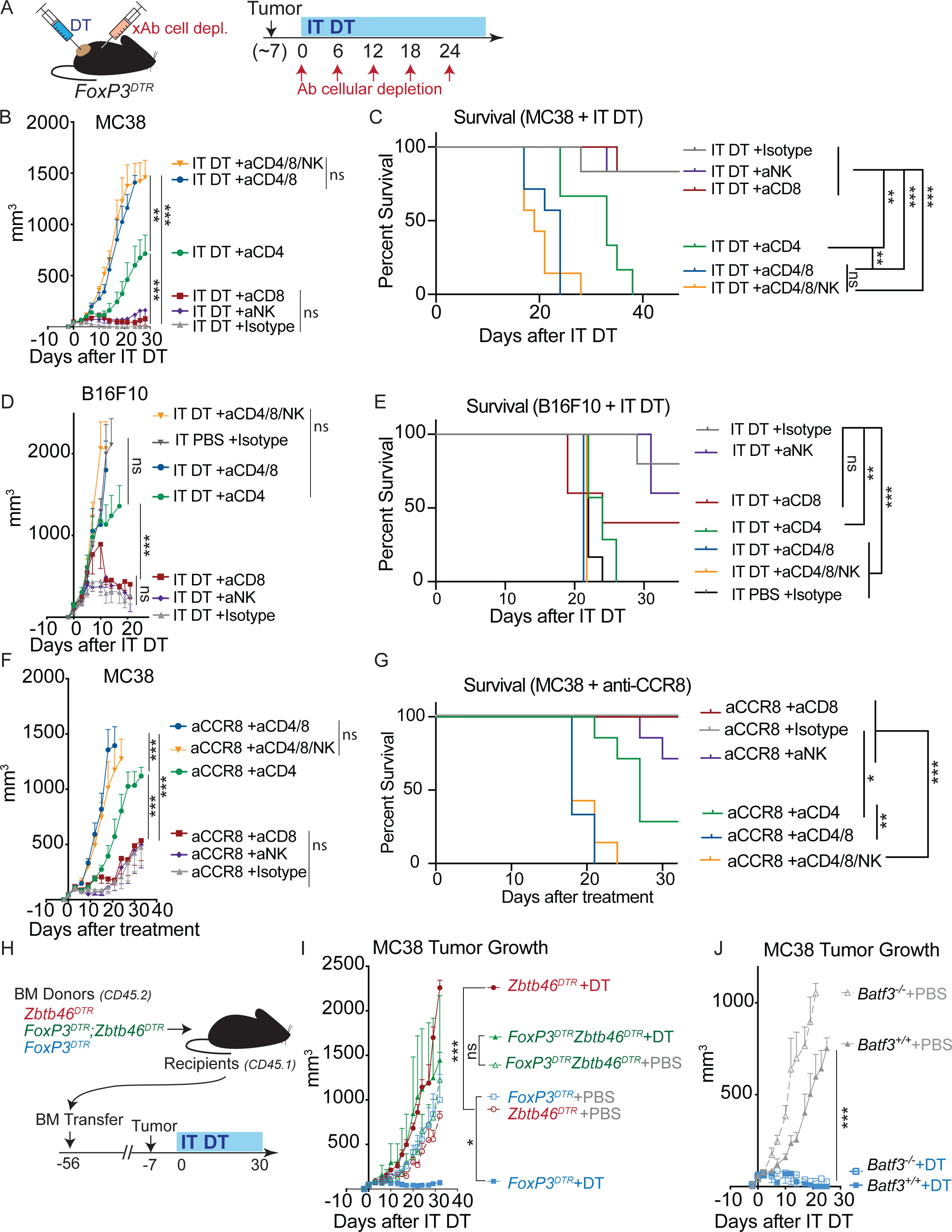
Conventional CD4+ T cells control tumors after IT Treg ablation. (A) Experimental design for cellular depletion studies in *Foxp3^DTR^* mice. IT Treg ablation was achieved through either IT DT or anti-CCR8 and began concurrent with cellular depletion. *In vivo* depletion antibodies were administered IP every 6 days for the duration of the study. (B) MC38 tumor growth in *Foxp3^DTR^* mice that received IT DT and indicated cellular depleting antibodies. (C) Animal survival after IT DT treatment and indicated cellular depletion. One experiment performed. (D) B16F10 tumor growth in *Foxp3^DTR^* mice that received IT DT and indicated cellular depleting antibodies. (E) Animal survival after IT DT and indicated cellular depletion. One experiment performed. (F) MC38 tumor growth after anti-CCR8 treatment and indicated cellular depletion. (G) Animal survival after anti-CCR8 and cellular depletion. One experiment performed. (H) Schematic of bone marrow chimera generation and treatment regimen in *Zbtb46^DTR^Foxp3^DTR^*chimeric mice. 250 ng IT DT and 80 ng IP DT was administered every 48-72 hours, once MC38 tumors reached 50mm^3^. (I) MC38 tumor growth in *Zbtb46^DTR^Foxp3^DTR^* chimeric mice. One experiment performed. (J) MC38 tumor growth in *Foxp3^DTR^Batf3^-/-^* mice. 250 ng IT DT was administered every 48-72 hours. Data are representative of 3 independent experiments. Data represent means ± SEM; *p < 0.05, **p < 0.01 and ***p < 0.001. B,D,F; I-J: Ordinary two-way ANOVA with Tukey’s multiple comparison test. C,E,J: Log-rank (Mantel-Cox) test. (n=5-7 mice/group).

This led us to test whether anti-CCR8 treatment, which is presumed to activate CD8^+^ T cell responses to tumors^28,33–37^, also exhibited similar cellular requirements for tumor control. With anti-CCR8 treatment, tumor control and survival were not prevented by NK or CD8^+^ T cell depletion alone, but were significantly reversed by depletion of CD4^+^ T cells, exactly as observed with the IT DT approach (Fig. 4F-G and Fig. S4C). Again, CD8^+^ T cells only participated in tumor control when CD4^+^ T cells were also depleted. Thus, despite the robust infiltration and activation of CD8^+^ T cells with IT T_reg_ ablation, CD8^+^ T cells do not appear to play a primary role in tumor control on their own. Overall, our findings instead highlight the largely underappreciated role for CD4^+^ T cells to control tumors without CD8^+^ T cells, particularly in the context of IT T_reg_ ablation.

As dendritic cells are integral to initiating and maintaining T cell responses, we next interrogated whether conventional DCs (cDCs) were required for tumor control elicited by IT T_reg_ ablation. We generated *Foxp3^DTR^;Zbtb46^DTR^* mice and then used their bone marrow to make bone marrow chimeric mice in which T_regs_ and cDCs could be inducibly ablated with DT administration^64,65^. Bone marrow chimera mice are required with *Zbtb46^DTR^* mice due to toxicity observed in intact *Zbtb46^DTR^*mice^64^. We implanted tumors into these mice and began IT DT treatment once tumors reached ∼50 mm^3^ (Fig. 4H). Tumor control with IT DT was prevented in mice that lacked cDCs (Fig. 4I). We also found that, in the absence of all cDCs, the increased frequency of conventional T cells within tumors that we had observed with IT T_reg_ ablation was prevented (Fig. S4D). These results suggest that cDCs were required for increasing the infiltration or maintenance of T cells within tumor tissues upon IT T_reg_ ablation.

As cDC1s are well-established as being integral to antitumor immunity in several contexts, such as checkpoint blockade immunotherapies, we next tested whether Batf3^+^, type I DCs (cDC1s) were required for tumor control with IT T_reg_ ablation^66–69^. We generated *FoxP3^DTR^;Batf3^-/-^* mice and then used IT DT treatment to ablate IT T_regs_ in mice that constitutively lacked cDC1s. We found that cDC1-deficient mice controlled tumors equivalently to control mice (Fig. 4J; Fig. S4E). These data indicate that cDC1s are not important for controlling tumors upon IT T_reg_ ablation and may also support the dispensability for CD8^+^ T cells previously observed. Notably, as a subset of cDC1s express CD8α, our previous experiments (Fig. 4A-G) selectively depleted CD8^+^ T cells using anti-CD8β depleting antibodies, thereby leaving this subset of cDC1s intact. These results highlight the dispensability of cDC1s in our model andsuggest that cDC2s, which were left intact in *Batf3^-/-^* mice, may be the critical subtype of cDC required for tumor control with IT T_reg_ ablation.

### Ablation of IT T_regs_ in a single tumor unleashes antitumor immunity against distal tumors

As disseminated metastatic cancers are in greatest need of treatment and would not be easily treated by IT delivery of agents to all tumors, we wanted to determine whether IT T_reg_ ablation in a single tumor would also boost systemic antitumor immune responses that could control untreated tumors. A precedent exists to support this hypothesis, as local radiation and IT injection of TLR agonists or checkpoint blockade inhibitors have been shown to not only impact the treated tumor, but distal tumors as well^46,48^. To test whether IT T_reg_ ablation alone in a single tumor also imparts control of distal tumors, we inoculated syngeneic tumors on both flanks of *FoxP3^DTR^* mice. We then initiated IT DT treatment in only one of the two tumors when the average volume of both tumors were ∼50 mm^3^. In mice with paired MC38 or B16F10 tumors, we found that IT DT treatment of one tumor in each mouse resulted in control of both the treated tumor and the untreated distal tumor (Fig. 5A,B and Fig. S5A-C). Control was only lost in distal tumors when DT treatment ceased due to the rejection of the primary treated tumors and our inability to continue IT DT treatments. Thus, IT T_reg_ ablation led to systemic control of a distal tumor even though distal tumors were not directly ablated of T_regs_. These results were indicative of the potential efficacy of IT T_reg_ ablation in a single tumor also controlling distant metastases.

**Figure 5.**
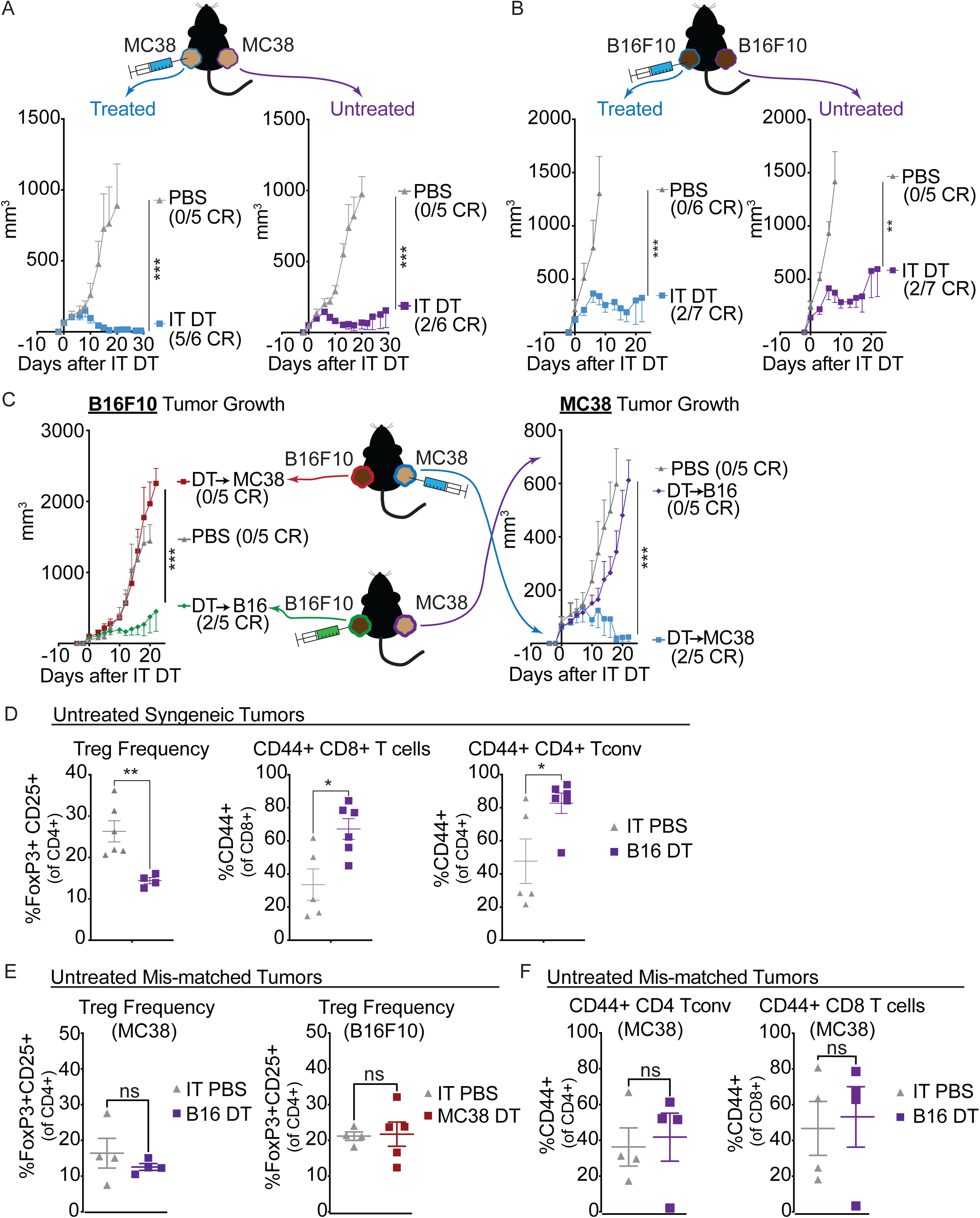
Ablation of IT Tregs in a single tumor unleashes antitumor immunity against syngeneic distal tumors. (A) MC38 tumor growth of an IT DT treated tumor and a distal, MC38 untreated tumor. (B) B16F10 tumor growth of an IT DT treated tumor and a distal, untreated B16F10 tumor. (C) B16F10 or MC38 tumor growth in *Foxp3^DTR^* mice bearing an MC38 tumor on the right flank and a B16F10 tumor on the left flank. B16F10 tumors are directly treated (green) or MC38 tumors are directly treated (blue). Data are representative of 5 independent experiments. (D) Quantification of Treg and activated CD4+ Tconv cell and CD8+ T cell frequencies in matched B16F10-tumor bearing mice. Tissue harvest performed at humane experimental endpoint for each cohort (D10 for PBS treated mice and D20 for IT DT treated mice). (E-F) Quantification of Treg frequencies or (F) activated CD4+ Tconv cells and CD8+ T cells in untreated MC38 or B16F10 tumors after IT DT in a mismatched B16F10 or MC38 tumor, respectively. Tissue harvest performed at humane experimental endpoint for each cohort (D20 for IT DT and IT PBS treated mice). Data are representative of 5 independent experiments for matched MC38 tumors and 2 independent experiments for matched B16F10 tumors. Data represent means ± SEM; *p < 0.05, **p < 0.01 and ***p < 0.001. A-C: Ordinary two-way ANOVA with Tukey’s multiple comparison test. D-F: Unpaired two-tailed T test. (n=4-7 mice/group).

We hypothesized that the control of the untreated distal tumor resulted from enhanced T cell activation against tumor antigens in the treated tumor that led to a stronger systemic T cell response against the specific tumor, which could target distant untreated tumors due to shared tumor antigens. However, distal tumor control could also be induced if IT T_reg_ ablation led to enhanced inflammation that promoted distinct T cell responses in the distant tumor. To discern which mechanism contributed to distal tumor control, we inoculated individual *FoxP3^DTR^* mice with two antigenically mismatched tumors, specifically MC38 on one flank and B16F10 on the other, and administered IT DT to only one tumor in each mouse (Fig. 5C). Unlike in mice with matched syngeneic tumors, in these mice with mismatched tumors, IT DT treatment only controlled the treated tumor and not the distal tumor (Fig. 5C). These results indicate that specific immunity generated within the treated tumor, likely reflecting T cell antigen specificity, contributed to the antitumor activity elicited against a distal, syngeneic tumor that was not directly treated. Furthermore, these results confirm that our IT DT approach results in local IT T_reg_ ablation that is restricted to the treated tumors and does not disrupt T_regs_ systemically, consistent with our finding that IT DT treatment does not induce systemic autoimmunity.

To investigate whether the T cell responses in distal untreated tumors were changed by IT T_reg_ ablation specifically with matched syngeneic tumors, we assessed the frequency and activation state of T cells in the tumors. In matched syngeneic tumors, there was a significant reduction of T_regs_ and an increase in activated (CD44^+^) CD8^+^ and CD4^+^ Tconv cells in the untreated tumors (Fig. 5D). We also observed an upregulation of PD1 and downregulation of CD62L on both T cell subsets in untreated tumors (Fig. S5D-E). However, in mismatched tumors, changes in T_reg_ frequencies and conventional T cell activation status were not observed, supporting the hypothesis that tumor antigen-specific T cell responses were only coordinated in the case of the matched tumors (Fig. 5E, F, Fig. S5F).

### Control of disseminated tumors is reversed by CD8^+^ or CD4^+^ T cell depletions

The impact of IT T_reg_ ablation on infiltration of CD4^+^ and CD8^+^ T cells into distant syngeneic tumors suggested that either or both of these populations may participate in distal tumor control. We therefore used cellular depletion studies to establish the immune cell dependencies for distal tumor control (Fig. 6A). As observed with single tumors, depletion of CD4^+^ T cells reversed the control of the IT T_reg_ ablated tumors as well as the matched distal tumors (Fig. 6B). Interestingly, however, while depleting CD8^+^ T cells alone did not impede the control of the primary treated tumors (as observed with single tumors, Fig. 4), control of the untreated distal tumor was largely reversed by CD8^+^ T cell depletion (Fig. 6B). These results demonstrate a context-specific dependency for CD8^+^ T cells in the control of primary versus disseminated cancer with IT T_reg_ ablation.

**Figure 6.**
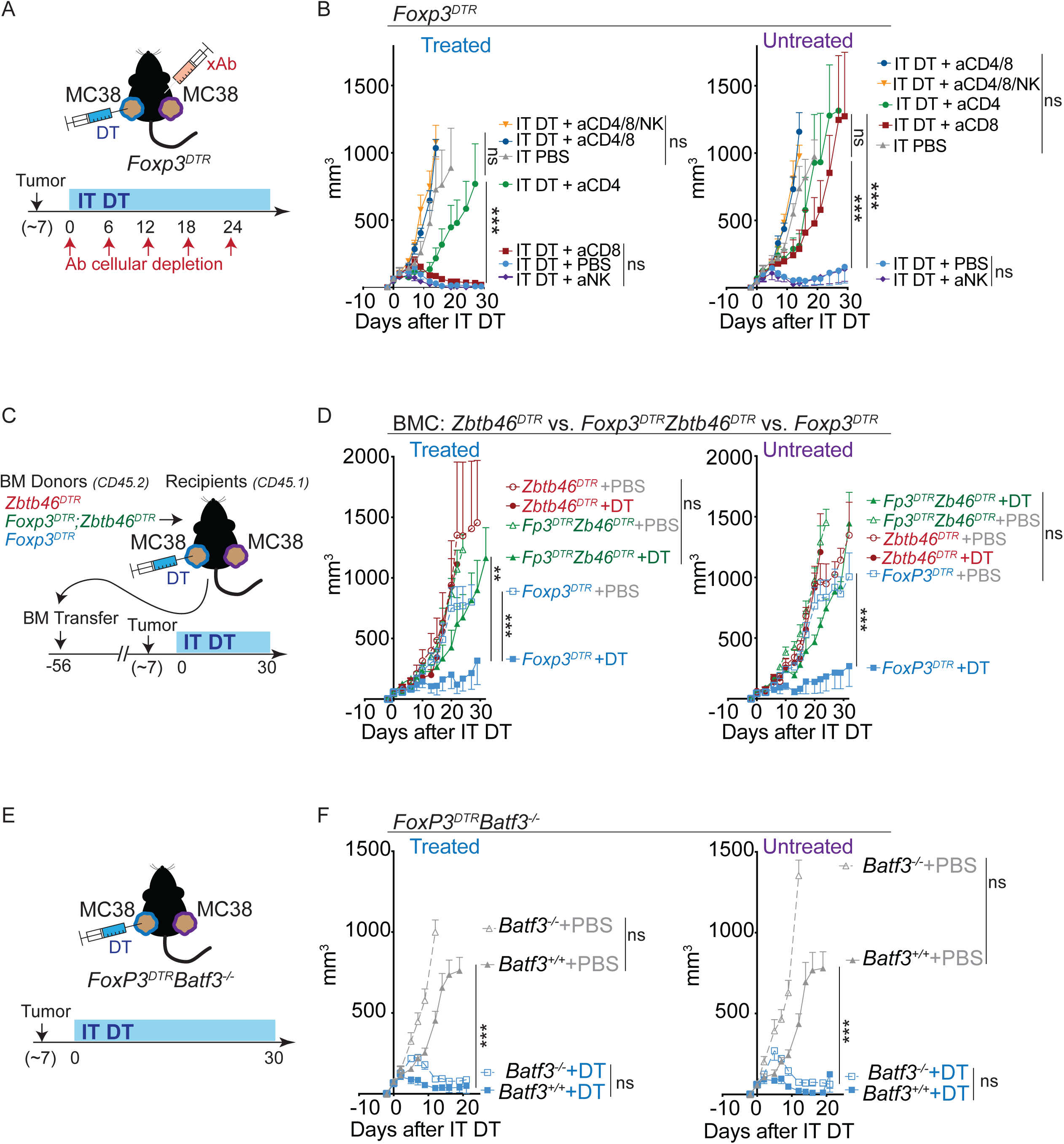
Control of disseminated tumors is reversed by CD8+ or CD4+ T cell depletions. (A) Schematic of tumor inoculation and cellular depletion regimen in *Foxp3^DTR^* mice. (B) MC38 tumor growth of an IT DT treated tumor or distal, untreated tumor. (C) Experimental schematic of *Zbtb46^DTR^Foxp3^DTR^* bone marrow chimera generation. (D) MC38 tumor growth in *Zbtb46^DTR^Foxp3^DTR^* bone marrow chimeric mice. One experiment performed. (E) Experimental setup of IT Treg depletion in *Foxp3^DTR^Batf3^-/-^* mice. (F) MC38 tumor growth in an IT DT treated tumor or a distal, untreated tumor in *Foxp3^DTR^Batf3^-/-^* mice. One experiment performed. Data are representative of 2-5 independent experiments. Data represent means ± SEM; *p < 0.05, **p < 0.01 and ***p < 0.001. B-G: Ordinary two-way ANOVA with Tukey’s multiple comparison test. (n=4-7 mice/group).

The requirement for CD8^+^ T cells to control matched disseminated tumors may indicate different cDC requirements for primary versus distal tumor control with IT T_reg_ ablation. Therefore, we again generated *Foxp3^DTR^;Zbtb46^DTR^* bone marrow chimera mice, implanted MC38 tumors on both flanks, and performed IT DT treatment of primary tumors (Fig. 6C). Conditional deletion of both IT T_regs_ and cDCs completely blocked the control of untreated distal tumors as well as the primary tumors (Fig. 6D). These data suggest that enhanced T cell activation observed after IT T_reg_ ablation is dependent upon the presence of cDCs. Given the requirement of CD8^+^ T cells in controlling tumor progression at a distal site, we expected that cDC1s might contribute to the clearance of dual tumors. However, *FoxP3^DTR^;Batf3^-/-^* mice with dual tumors and IT DT treatment of a primary tumor showed no impairment in the control of either tumor (Fig. 6E, F). Thus, cDCs that do not require Batf3 function were sufficient to promote tumor control with IT T_reg_ ablation, suggesting again that cDC2s may act as the critical mediators of primary and distal tumor control.

### IT T_reg_ ablation enhances tumor antigen acquisition by cDC2s

To investigate how IT T_reg_ ablation impacted cDCs within the tumor, we implanted MC38 tumor cells that expressed an mCherry fluorescent protein as a trackable tumor antigen^67^. Additionally, we pre-depleted CD8^+^ T cells in *FoxP3^DTR^*mice prior to tumor implantation to reduce any complications due to CD8^+^ T cell-mediated cytotoxicity in enhancing tumor antigen release. When tumors reached ∼50 mm^3^, we administered 3 doses of IT DT every other day and at day seven, tumors and tdLNs were analyzed by flow cytometry to quantify immune cells that had phagocytosed fluorescent tumor antigen, in particular cDC1s and cDC2s (Fig. 7A). We found a significant increase in the frequency of mCherry^+^ CD11c^+^MHC-II^+^ DCs in the tumors but not tdLN of IT T_reg_ ablated mice (Fig. 7B and Fig. S6A). Interestingly, by further gating on CD103^+^ cDC1s and CD11b^+^ cDC2s, we observed that only the cDC2s exhibited significantly increased tumor antigen uptake in tumors upon IT T_reg_ ablation (Fig. 7C and Fig. S6B). In addition, as observed previously with systemic T_reg_ depletion^16,70–72^, we observed increased expression of CD80 and CD86 costimulatory ligands on CD11c^+^MHC-II^+^ DCs, especially on DCs that were actively phagocytosing tumor antigen (i.e. mCherry^+^) (Fig. S6C-E). Interestingly, however, we again observed that only the tumor antigen+ cDC2s, and not antigen+ cDC1s, exhibited increased expression of CD80 and CD86 ligands in tumors (Fig. 7D). This highlights a crucial role for IT T_regs_ in limiting cDC2 tumor antigen uptake and maturation in tumors, which likely leads to reduced antitumor CD4^+^ and CD8^+^ antitumor T cell responses.

**Figure 7.**
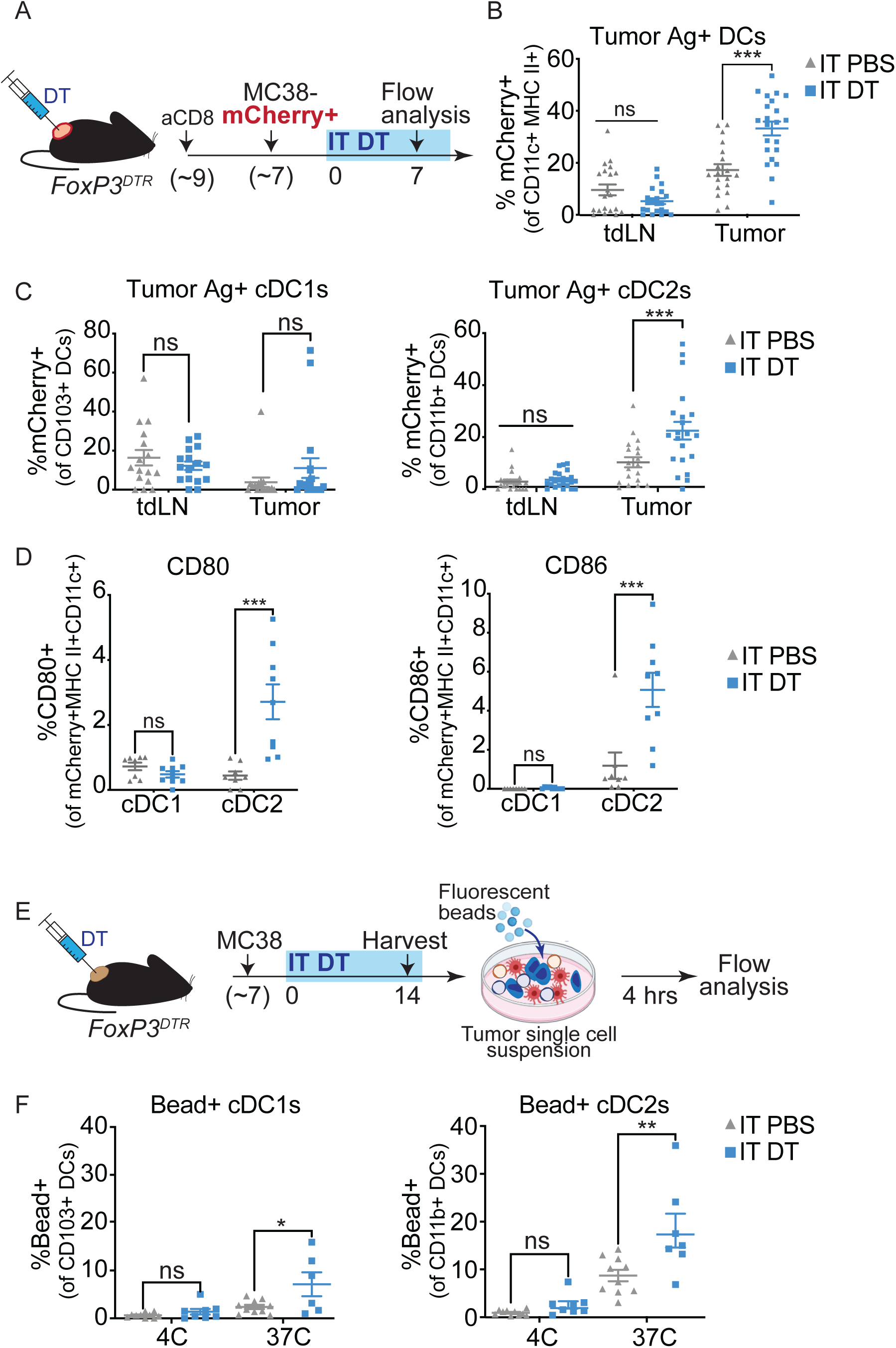
IT Treg ablation enhances tumor antigen acquisition by cDC2s. (A)Experimental schematic. *Foxp3^DTR^* mice received anti-CD8 depleting antibodies two days prior to mCherry+ MC38 tumor inoculation and every six days thereafter. IT DT began when tumors reached 50mm^3^. (B) Quantification of mCherry+ DCs after IT PBS or IT DT treatment. (C) Quantification of mCherry+ cDC1s or cDC2s in tumor and tdLNs. (D) Quantification of CD80 and CD86 expression on mCherry+ cDC1s and mCherry+ cDC2s. (E) Schematic of experimental approach in *ex vivo* assays. *Foxp3^DTR^*mice were inoculated with MC38 tumors and dosed with IT DT when tumors reached 50mm^3^. Tumors were harvested and plated with fluorescent beads after seven days of IT DT treatments. (F) Frequencies of cDC1s or cDC2s that phagocytosed beads at either 4C or 37C. *In vivo* data are representative of 2 independent experiments. *Ex vivo* data are representative of 3 independent experiments. Data represent means ± SEM; *p < 0.05, **p < 0.01 and ***p < 0.001. B-E; G,H: Ordinary two-way ANOVA with Tukey’s multiple comparison test. (n=4-7 mice/group).

To directly test whether cDC2s from tumors exhibited increased phagocytic capacity after IT T_reg_ ablation, we tested tumor single cell suspensions of tumors harvested from IT DT versus IT PBS treated mice for uptake of fluorescent beads *ex vivo* (Fig. 7E). In this assay, we again found that the primary subset of cDCs that had increased phagocytosis of bead antigen in the absence of IT T_regs_ were cDC2s, although IT T_reg_ ablation also enhanced bead uptake in cDC1s, but to a lesser extent (Fig. 7F). These data reveal a novel mechanism by which IT T_regs_ restrict antitumor T cell responses, by restricting tumor antigen uptake locally within tumors. Here, we demonstrate that IT T_regs_ restrict cDC2 uptake of tumor antigen within tumor tissue, thereby dampening CD4^+^ Tconv cell responses against both local and disseminated tumors. By eliminating T_regs_ specifically within tumors, we found that cDC2 activation and phagocytic capacity were dramatically enhanced, unleashing a CD4^+^ Tconv cell response that acts independently of cDC1s to control tumors. Thus, intratumoral T_reg_ ablation activates an alternative mechanism of tumor control that depends on cDC2s and CD4^+^ Tconv cells.

## DISCUSSION

Although modulating T_reg_ function is a key approach to boosting cancer immunotherapy, the role of intratumoral T_regs_ and the mechanisms by which they suppress anti-tumor T cell responses remains incompletely defined. To address this gap in knowledge, we leveraged the *FoxP3^DTR^* mouse coupled with low-dose IT DT treatment to selectively ablate T_regs_ in tumors. These experiments revealed that T_reg_ ablation within tumor tissue itself was sufficient to limit tumor growth. Importantly, this tumor control occurred without any autoimmune side effects, which are common with T_reg_-targeting approaches. Removal of IT T_regs_ increased conventional CD4^+^ and CD8^+^ T cell activation in tumors. However, CD4^+^ Tconv cells and conventional type II dendritic cells (cDC2s) were the primary cells responsible for tumor control with IT T_reg_ ablation. These cellular requirements extended to clinical cancer interventions, as we also observed a requirement for CD4^+^ Tconv cells, rather than CD8^+^ T cells, after anti-CCR8 antibody-mediated IT T_reg_ depletion. These data are consequential, as many combinatorial therapeutics aimed at reducing T_reg_ suppression are thought to act by bolstering antitumor CD8^+^ T cell responses^15,24,28,53,60,61^. For example, two recent studies specifically disrupted IT T_regs_, even with anti-CCR8 antibodies, and showed increased frequencies of total and tumor-specific CD8+ T cells in tumors, as we also observed in this study, but did not functionally validate their importance by cellular depletion experiments^27,38^. Still other groups have disrupted intratumoral T_reg_ function through either targeting tumor-specific T_reg_ genes or disrupting their recruitment, and in these cases the mechanisms of tumor control were shown to be driven by CD8^+^ T cell responses^63,67,73^. Here we show, by CD8^+^ T cell depletion, that CD8^+^ T cells are not required for tumor control, as previously hypothesized, and instead reveal the critical importance of CD4^+^ Tconv cells for tumor control upon total IT T_reg_ ablation.

Here, we also established that systemic responses against disseminated cancers, e.g. metastases, is enhanced by IT T_reg_ ablation in a single tumor. Enhanced antitumor responses only occurred when we disrupted IT T_reg_ suppression in a tumor of the same origin as an untreated distal tumor. However, in this setting of disseminated cancer, we found that both CD8^+^ and CD4^+^ Tconv cells were essential for controlling untreated distal tumors. These findings align with other studies that demonstrated an abscopal therapeutic effect after local IT immunotherapies^46,47,60^, but extend such findings by revealing the importance of cDC2s and CD4^+^ Tconv cells in the control of distal and untreated tumors. Further, our findings are unique in showing that systemic immune responses can be elicited by antibody-independent, non-Fc receptor activating, depletion of IT T_regs_.

By disrupting conventional DCs, we also uncovered a critical role for cDCs in promoting T cell responses against tumors. Most cancer immunotherapeutic studies have focused on the role of cDC1s in promoting T cell responses to tumors^74–77^. As such, efforts have focused on targeting the cDC1-CD8 T cell axis in human cancer immunotherapies, such as by selectively disrupting T_reg_ interactions with cDC1s^67^. However, as IT T_reg_ ablated tumors appear to require CD4^+^ Tconv cell responses, it is possible that a different subset of conventional DCs are important for tumor control in this setting. Indeed, while CXCR3+ T_regs_ were shown to selectively impede CD8^+^ T cell responses against tumors, our data demonstrate that removal of all IT T_regs_ induces tumor regression through a cDC2, and cDC1-independent, mechanism. Recent literature has demonstrated that mature DCs enriched in immunoregulatory molecules (mregDCs) can recruit T_regs_, allowing for their accumulation around lymphatic vessels in the peripheral tumor stroma^78^. By disrupting this mregDC-T_reg_ axis, tumor immunosurveillance was enhanced^79^. While our data suggest a prominent role for cDC2s in stimulating anti-tumor immune activity, it remains possible that mregDCs may also contribute to tumor rejection in the absence of T_regs_. Together, these findings are consequential, as they represent a parallel arm of antitumor T cell immune function that may bolster tumor control in cDC1 or CD8^+^ T cell non-responsive tumors.

We also show that IT T_regs_ blocked the acquisition of tumor antigens by cDC2s within tumor tissue. In the absence of all cDCs, we abrogated enhanced T cell activation and expansion induced by IT T_reg_ ablation. Recently, Binnewies et. al. demonstrated a critical role for cDC2 responses in the tdLN after systemic T_reg_ removal^17^. These studies demonstrated that in the absence of T_regs_, cDC2s drove stronger CD4^+^ T cell responses by enhancing the trafficking of cDCs presenting tumor-derived antigens to CD4^+^ Tconv cells in the tdLN^17,80^. Furthermore, a correlation between the abundance of IT cDC2s and the accumulation of ITCD4^+^ Tconv cells was demonstrated in the absence of T ^68^. However, whereas previous studies have suggested that the migration of cDC2s is regulated by T_regs_, here we show that the uptake of tumor antigens is also controlled by IT T_regs_, and that regulation by cDC2s is important directly within tumors. That is because, unlike others^17^, we found that FTY720 treatment did not impede tumor control with IT T_reg_ ablation, strongly suggesting that increased priming of CD4^+^ T cells in tdLNs is not required for the efficacy of IT T_reg_ ablation. This may be due to the impact of IT-specific Treg ablation performed here, as opposed to systemic T_reg_ ablation, and deserves further investigation. In addition, it is noteworthy that IT T_reg_ ablation was beneficial compared to systemic T_reg_ ablation because it was sufficient to promote tumor control without causing autoimmune pathology commonly observed in other studies.

Maintenance of immune homeostasis is a vital function of T_regs_^7,13^. Thus, disrupting global T_reg_ function incurs severe autoimmune disease in humans and in mice^11,12,14^. This motivated our efforts to locally deplete T_regs_ only in tumors to see if this was sufficient for tumor control and would retain critical T_reg_ suppressive functions in peripheral tissues. While others have employed similar approaches to disrupt tdLN and tumor T_regs_ simultaneously^27^, or use anti-CTLA4 targeting antibodies to disrupt IT T_reg_ function^43^, our approach remains distinct as we only target IT-T_regs_ without the use of Fc-engaging antibodies and show tumor control likely occurred without tdLN T cell priming by use of FTY720. However, even though we did not detect changes in CD4^+^ Tconv or CD8^+^ T effector function within the tdLN, and tumor control was retained even with FTY720 treatment, it remains possible that low-level depletion of tdLN T_regs_ also contributed to antitumor immunity. However, our data that show a loss of tumor control when distant tumors are mismatched from the treated tumor does suggest that IT T_reg_ ablation acts locally within tumors. Thus, our results support the feasibility of strategies that selectively disrupt IT T_regs_ for safe and effective cancer immunotherapy.

Current clinical data demonstrate poor prognostic outcomes in tumors that are cDC1 or CD8^+^ T cell low^81,82^. These tumors have been shown to respond poorly to anti-PD-1 checkpoint blockade as well as other immunotherapies^77,83,84^. In these cases, failed tumor immunosurveillance was attributed to the downregulation of MHC I molecules on the tumor cell surface, or the absence of tumor neoantigens, capable of eliciting CD8 T cell responses^85–88^. Our data identify a novel mechanism to combat CD8-resistant tumors, or synergize with such existing therapeutic strategies, by unleashing cDC2s and CD4^+^ Tconv cell-mediated tumor control by IT T_reg_ ablation.

In summary, here we show that IT T_reg_ ablation is sufficient to promote tumor control and uncover a novel function of IT T_reg_ function - to suppress cDC tumor antigen uptake. IT T_reg_ function suppresses effective CD4^+^ Tconv cell responses locally in tumors and disrupts the ability of both CD4^+^ and CD8^+^ T cells to cooperate in the control of distant untreated tumors. Our findings also extend to a clinically viable mode of IT T_reg_ depletion using an anti-CCR8 antibody delivered systemically. These data are consequential, as they demonstrate a novel arm of adaptive immunity that can be leveraged to enhance the immunotherapeutic treatment of patients with cancer.

## Methods

All of the experiments were conducted according to the Institutional Animal Care and Use Committee guidelines of the University of California, Berkeley. Further information on the committees that approved protocols used in this publication can be found in each corresponding section.

### *In vivo* animal studies

Mice used were bred onto a C57BL/6 background for a minimum of ten generations. Littermates or age-matched control mice (8-20 weeks old) were used in all mouse experiments. *Foxp3^GFP-DTR^*mice express human diphtheria toxin receptor and EGFP genes from the Foxp3 locus without disrupting expression of the endogenous FoxP3 gene^11^. These mice were a gift from Dr. Rudensky (Memorial Sloan Kettering Institute; JAX:016958). *Foxp3^GFP-DTR^;Kras*^G12D^*;p53*^fl/fl^ mice were kindly provided by Dr. Jacks (Massachusetts Institute of Technology). *Zbtb46^DTR^* mice were originally characterized by Dr. Nussenzweig (Rockefeller University) and *Batf3^-/-^* mice were characterized by Dr. Murphy (Washington University School of Medicine). Both *Zbtb46^DTR^* mice and *Batf3^-/-^* mice were obtained from Jackson laboratories (JAX:019506; and JAX:013755, respectively) and backcrossed with *Foxp3^GFP-DTR^* mice in our laboratory. For diphtheria toxin treatments, mice were injected with 1 μg intraperitoneal (IP) or 250 ng intratumoral (IT) every other day up to three times per week. For syngeneic tumor studies, C57BL/6 mice were subcutaneously inoculated in 100uL of PBS with 5x10^5^ MC38 cells or 5x10^5^ B16F10 cells. For autochthonous tumor studies, sarcomas were induced in *Foxp3^GFP-DTR^;Kras*^G12D^*;p53*^fl/fl^ mice by intramuscular (IM) injection of the hind calf muscle with replication-incompetent lentiviruses express Cre recombinase as reported previously. Mice were monitored three times per week for palpable sarcoma formation beginning 50 days after IM injection. Tumor measurements were performed blindly using calipers three times per week. A single operator measured each tumor in three dimensions for each tumor measurement. All experiments were performed in accordance with the Institutional Laboratory Animal Care and Use Committee (IACUC) and the Office of Laboratory Animal Care (OLAC) at the University of California, Berkeley.

### Lentiviral production

293T cells were transfected with d8.2 (gag/pol), CMV-VSV-G and a transfer vector expressing Cre to produce lentivirus.

### Cell lines

MC38 and B16F10 cell lines were kindly provided by Dr. Bluestone (UCSF). All MC38 cell lines expressed luciferase with or without co-expression of mCherry. All cell lines were maintained in DMEM (GIBCO) supplemented with 10% FBS, sodium pyruvate (GIBCO), 10mM HEPES (GIBCO), and penicillin-streptomycin (GIBCO). Tumor cells were grown at 37C with 5% CO_2._

### Murine Lymphocyte isolation

Single cell suspensions were generated from lymphoid organs through mechanical disruption of harvested tissues and were passed through 40um filters. Suspensions were resuspended in ice-cold FACS buffer (PBS supplemented with 2mM EDTA and 2% FBS). Spleens were subjected to red blood cell lysis using ACK lysis buffer (150mM NH4Cl, 10mM KHCO3, 0.1mM NA2EDTA, pH 7.3). Resected tumors were minced to 1mm3 isolates and digested in RPMI media supplemented with 4-(2-hydroxyethyl)-1-piperazineethanesulfonic acid (HEPES), 20mg/mL DNase I (Roche), and 125U/mL collagenase D (Roche). Tumor isolates were digested in an orbital shaker for 45 minutes at 37C. All cell suspensions were passed through 40um filters before cell staining or in vitro stimulation. Cytokine staining was performed on 3-5x10^6^ cells, after 210 minutes of *in vitro* stimulation in Opti-MEM media supplemented with Brefeldin A (eBioscience), 10ng/mL phorbol 12-myristate 13-acetate (PMA)(Sigma), and 0.25uM ionomycin (Sigma). Fixation and permeabilization of cells was performed for intracellular staining using the Tonbo Foxp3 Transcription Factor Staining Buffer Kit. In some experiments, e.g. staining of APCs for in vivo acquisition of mCherry tumor antigen, cells were fixed with 4% PFA on ice for 15 minutes.

### Flow cytometry

Flow cytometry was performed on a BD LSR Fortessa X20 (BD Biosciences) and datasets were analyzed using FlowJo software (Tree Star). All samples analyzed were reconstituted in single cell suspensions using ice-cold FACS buffer (PBS with 2mM EDTA and 2% FBS). Dead cells were stained with Live/Dead Fixable Blue or Aqua Dead Cell Stain kit (Molecular Probes) in PBS for 20 minutes at 4C. Cell surface proteins were stained for 20 minutes at 4C using a mixture of fluorophore-conjugated antibodies. Surface marker stains for murine samples were carried out with anti-mouse CD4 (RM4-5, BioLegend), anti-mouse CD8a (53-6.7, BioLegend), anti-CD25 (PC61, BioLegend), anti-mouse CD44 (IM7, BioLeged), anti-mouse CD45 (30-F11, BioLegend), anti-mouse CD103 (2E7, BioLegend), anti-mouse PD1 (29F.1A12, BioLegend), anti-mouse CCR8 (SA214G2, Biolegend), anti-mouse MHCII (M5/114.15.2, BioLegend), anti-H-2Kb MuLV p15E Tetramer-KSPWFTTL (NIH tetramer core), anti-mouse TIGIT (Vstm3, Biolegend), anti-mouse ICOS (C398.4A, Biolegend), anti-mouse CD62L (MEL1-14, Biolegend) anti-mouse CD11b (M1/70, Biolegend), anti-mouse CD3e (17A2, Biolegend), anti-mouse B220 (FA3-6B2, Biolegend), anti-mouse F4/80 (BM8, Biolegend), anti-mouse CD80 (M16-10A1, Biolegend), anti-mouse CD86 (GL-1, Biolegend) and anti-mouse PD-L1 (10F.9G2, Biolegend). anti-mouse CD11c (N418, Biolegend) in PBS for 20 minutes at 4C. Cells were fixed using the Tonbo Foxp3 Transcription Factor Staining Buffer Kit, prior to intracellular staining. Intracellular staining was performed using anti-mouse anti-mouse Foxp3 (FJK-16S, eBioscience), anti-mouse IL-2 (JES6-5H4, Biolegend), anti-mouse TNF-a (MP6-XT22, BioLegend), anti-mouse IL-10 (JES5-16E3, Biolegend), anti-mouse IFNg (XMG1.2, eBioscience) in PBS for 1 hour at 4C, according to manufacturer’s instructions. Cells were resuspended in FACS buffer and filtered through a 40-mm nylon mesh before data acquisition.

### In vivo antibody-mediated cell depletion

Immune cell depletion of CD8+ T cells, CD4+ T cells or NK cells was achieved through intraperitoneal injection of 200ug per mouse of each respective depleting antibody. Antibody-based depletion was started at the time of IT DT and was readministered every 6 days throughout the study. CD8b.2 (Clone 53-5.8, Leinco Technologies, Catalog # C2832) was used for CD8+ T cell depletions. Clone GK1.4 (Leinco Technologies, Catalog #C1333) was used for CD4+ T cell depletions. Clone PK136 (Leinco Technologies, Catalog #N123) was used for NK cell depletions. Anti-CCR8 mIgG2a (Gilead Sciences) was administered at 200 ug IP seven days after tumor inoculation and was re-dosed every 3 days.

### Bone marrow chimera generation

Host CD45.1+ C57/BL6 mice were administered two doses of 5 Gy irradiation, 16 hours apart. An Xrad320 (Precision X-Ray Irradiation) was used to administer irradiation. 4 hours after the second irradiation treatment, 10x106 bone marrow cells were transferred to host mice via tail vein injection. Bone marrow was harvested from donor *Zbtb46^DTR^*, *Foxp3^DTR^*, or *Zbtb46^DTR^Foxp3^DTR^* CD45.2+ C57/BL6 mice. Femurs and tibias were harvested from euthanized donor mice and flushed with ice-cold RPMI supplemented with 2% FBS, using a 27-gauge syringe. Red blood cells were removed from bone marrow preparations by using ACK lysis buffer (150mM NH4Cl, 10mM KHCO3, 0.1mM NA2EDTA, pH 7.3). Bone marrow was then washed with ice-cold PBS three times prior to tail vein injection. After bone marrow transfer, mice were administered water supplemented with sulfamethoxazole-trimethoprim oral suspension (Ani Pharmaceuticals) for 30 days and were monitored daily. Bone marrow was permitted to reconstitute recipient mice for 60 days. Following reconstitution, 5x10^5^ MC38 tumor cells were subcutaneously inoculated. 7 days after tumor inoculation, mice were administered either IT PBS or a combination of 250 ng IT DT and 80 ng intraperitoneal DT.

### Statistical methods

p values were obtained from unpaired two-tailed Student’s t tests for all statistical comparisons between two groups, and data were displayed as mean ± SEMs. For multiple comparisons, one-way ANOVA was used. For tumor growth curves, two-way ANOVA was used with Tukey’s multiple comparisons test performed at each time point or by multiple regression analysis. Survival analysis was performed using one-way chi-square test or Log-rank (Mantel-Cox) test. p values are denoted in figures by *p < 0.05, **p < 0.01, and ***p < 0.001.

## Data availability

Values Data supporting the findings of this study are available within the article and supplementary files or from the corresponding authors upon request. Source data are provided with this paper.

## Acknowledgements

We thank David Raulet, Ellen Robey, Greg Barton and members of the DuPage laboratory for reviewing the manuscript and providing feedback. We also thank Hector Nolla, Alma Valleros, Kartoosh Heydari, Melaine Delcroix and Harman Dhaliwal of the UC Berkeley Cancer Research Laboratory Flow Cytometry Facility. Furthermore, we thank all the members of the DuPage lab for providing guidance on the research approach. M.D is a Pew-Stewart Scholar and a St. Baldrick’s Scholar with generous support from Hope with Hazel.

## Author Contributions

Conceptualization and methodology, M.D. and A.B.; Investigation, A.B., B.G. and M.D.; writing - original draft, M.D. and A.B; writing - review & editing, M.D., A.B., B.G., C.Z.; supervision, M.D. and A.B.; funding acquisition, M.D.

## Additional information

**Correspondence and requests for materials** should be addressed to Alissa Bockman or Michel DuPage.

