## Supplementary Data for "Intratumoral Treg ablation is sufficient to mediate tumor control systemically without autoimmunity"

### SUPPLEMENTAL FIGURE 1

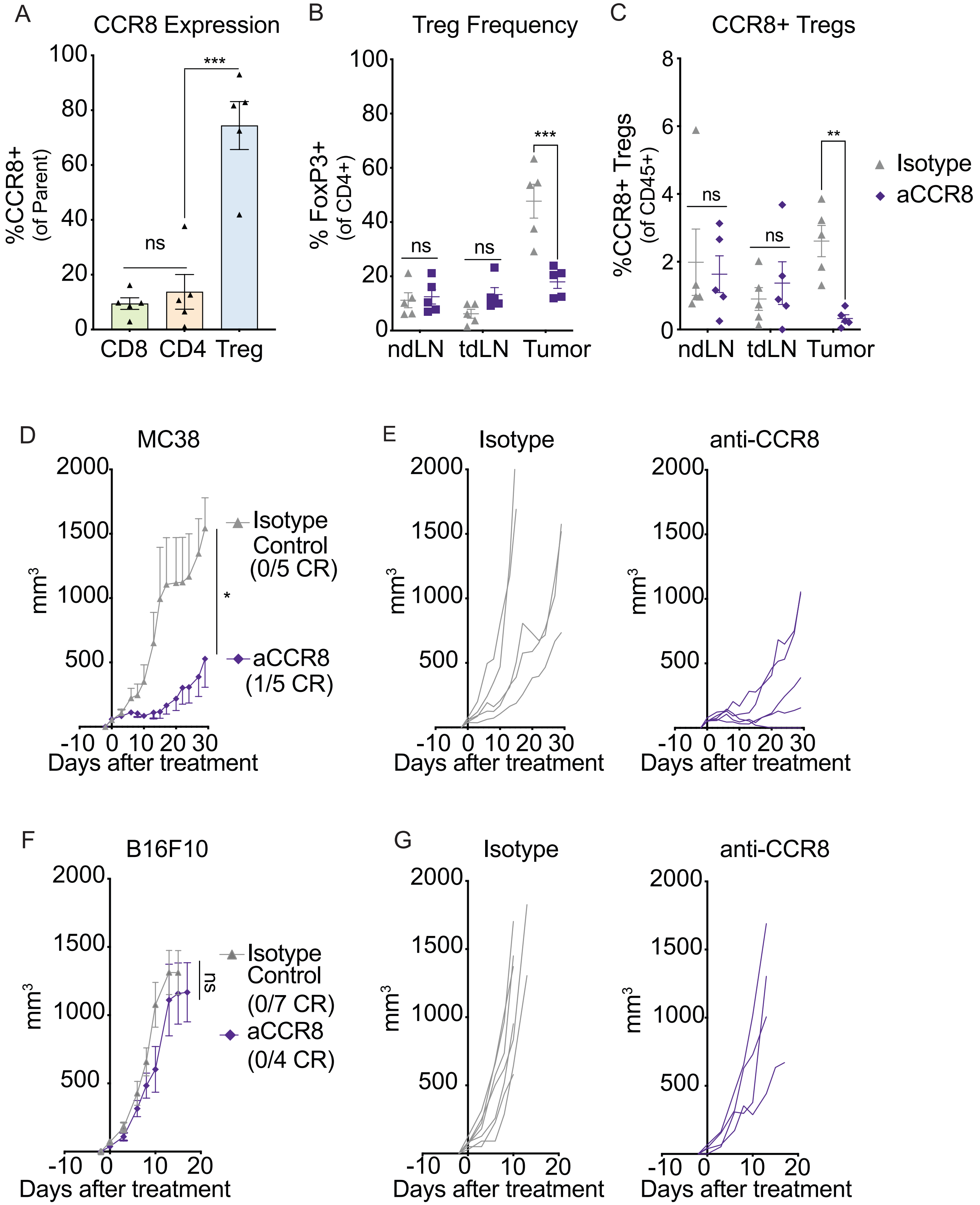

**Supplemental Figure 1. Intratumoral Treg depletion is sufficient to control tumor growth.**
(A) Quantification of CCR8 expression on MC38 tumor-infiltrating CD8+ T cells, CD4+ Tconv cells and Tregs.
(B) Frequencies of FoxP3+ Tregs in ndLN, tdLN and tumor tissues, calculated as a proportion of CD4+ lymphocytes.
(C) Quantification of CCR8+ Tregs after anti-IgG2A isotype or anti-CCR8 treatment in tdLN, ndLN and tumor tissue.
(D) MC38 tumor growth after anti-IgG2A isotype or anti-CCR8 treatment.
(E) Individual MC38 tumor growth curves of anti-IgG2A isotype or anti-CCR8 treated mice.
(F) B16F10 tumor growth after anti-IgG2A isotype or anti-CCR8 treatment.
(G) Individual B16F10 tumor growth curves of anti-IgG2A isotype or anti-CCR8 treated mice.
Data are representative of 2 independent experiments. Data represent means ± SEM; $*p < 0.05$, $**p < 0.01$ and $***p < 0.001$. A: Unpaired two-tailed T test. B-C;D;F: Ordinary two-way ANOVA with Tukey's multiple comparison test. (n=5-7 mice/group).

SUPPLEMENTAL FIGURE 2

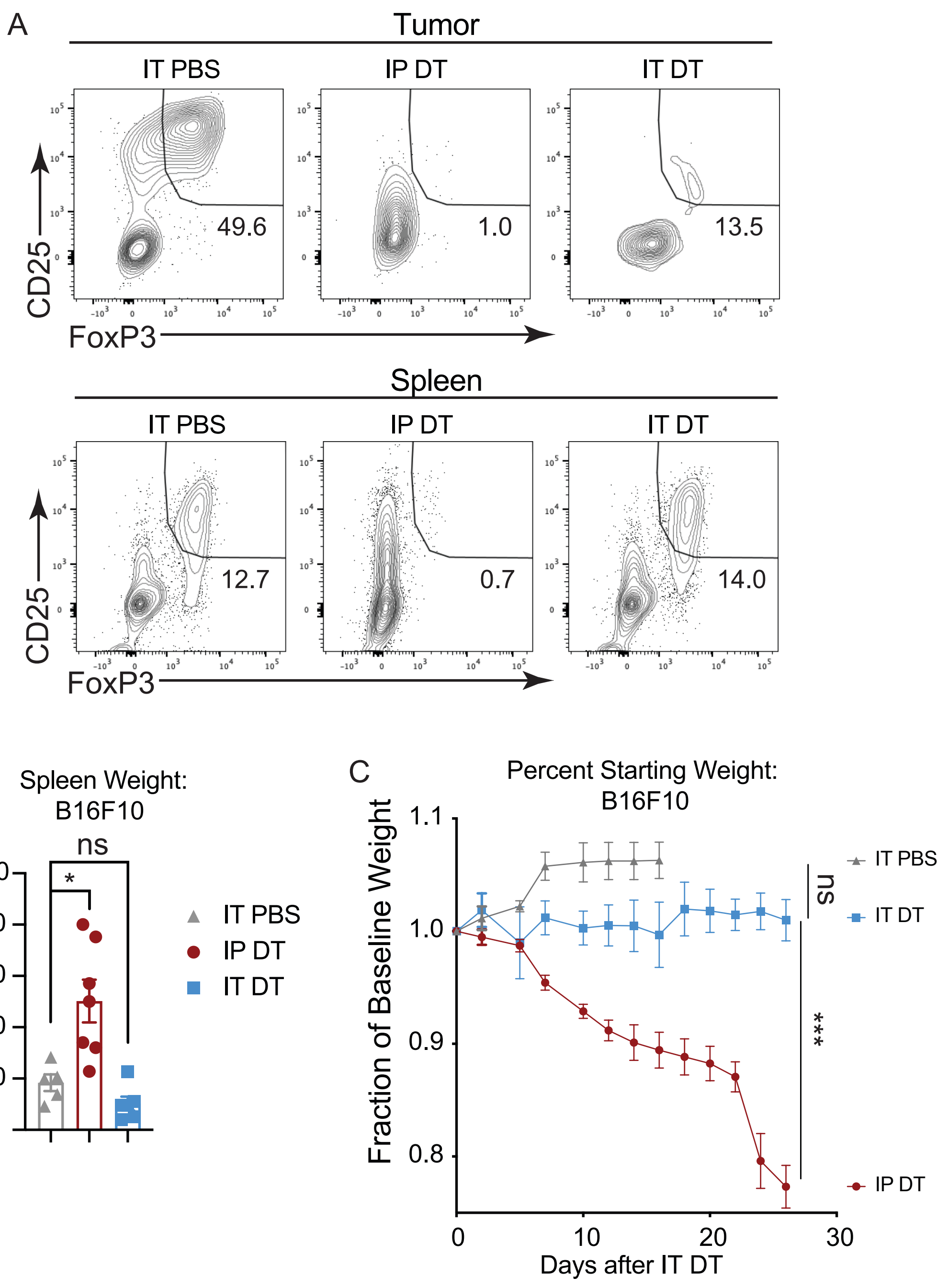


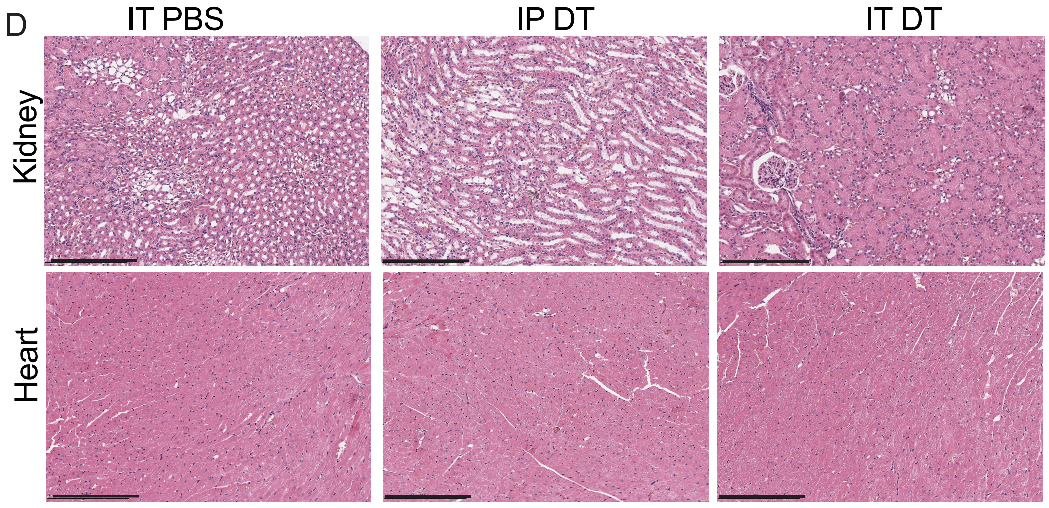


E **Pathology score:** severely damaged (++), damaged (+), normal (-)

| Group | Sample ID | Heart | Kidney | Treatment |
|---|---|---|---|---|
| 1 | 494 | - | + | IT PBS |
| | 495 | - | + | |
| 2 | 238 | - | + | IP DT |
| | 386 | - | ++ | |
| 3 | 239 | - | - | IT DT |
| | 385 | - | - | |

SUPPLEMENTAL FIGURE 2 (cont.)

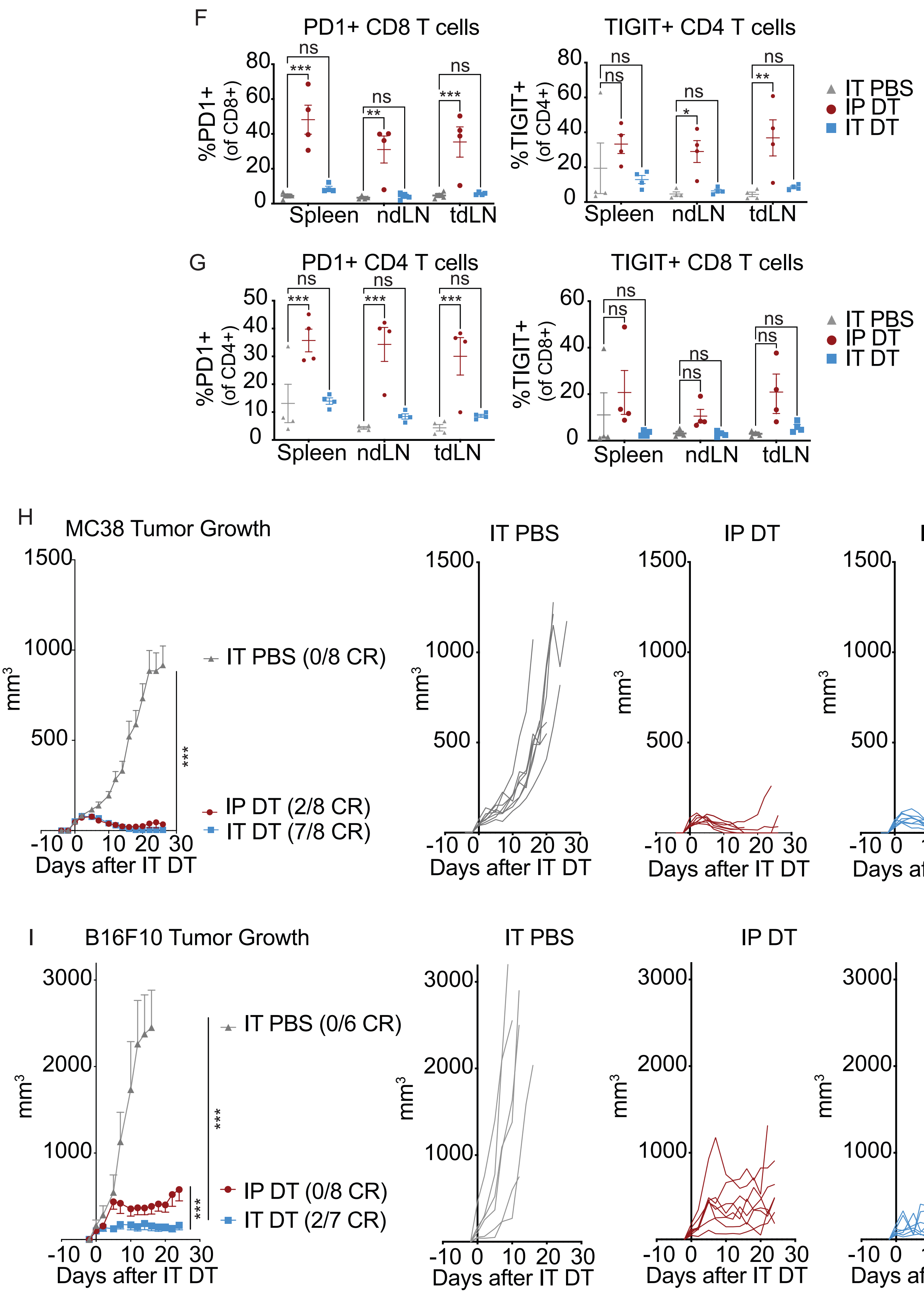

**Supplemental Figure 2. IT Treg ablation, unlike systemic Treg ablation, does not cause autoimmune pathology.**

(A) Representative flow cytometry plots of CD25 vs. FoxP3 Treg populations in MC38 tumors (top) or spleens (bottom) after IT PBS, IP DT or IT DT treatment in *Foxp3*$^{DTR}$ mice.

(B) Spleen weight in B16F10 tumor-bearing *Foxp3*$^{DTR}$ mice.

(C) Body weight of B16F10 tumor-bearing *Foxp3*$^{DTR}$ mice after IT PBS, IP DT or IT DT treatment.

(D) Representative images depicting Hematoxylin and eosin staining of kidney and heart (scale bar = 200um) tissues after IT PBS, IP DT or IT DT treatment in MC38 tumor-bearing *Foxp3*$^{DTR}$ mice. Images are representative of two biological replicates for each treatment group.

(E) Pathology scores of IT PBS, IP DT or IT DT treated mice.

(F) Frequency of PD1+ or TIGIT+ CD8+ T cells.

(G) Frequency of PD1+ or TIGIT+ CD4+ Tconv cells.

(H-I) MC38 or (I)B16F10 tumor growth and spider plots of individual tumor growth in IT PBS, IP DT or IT DT treated mice.

Data are representative of 2 independent experiments. Data represent means ± SEM; *p < 0.05, **p < 0.01 and ***p < 0.001. B: Unpaired two-tailed T test. C;F-I: Ordinary two-way ANOVA with Tukey's multiple comparison test. (n=4-7 mice/group).

SUPPLEMENTAL FIGURE 3

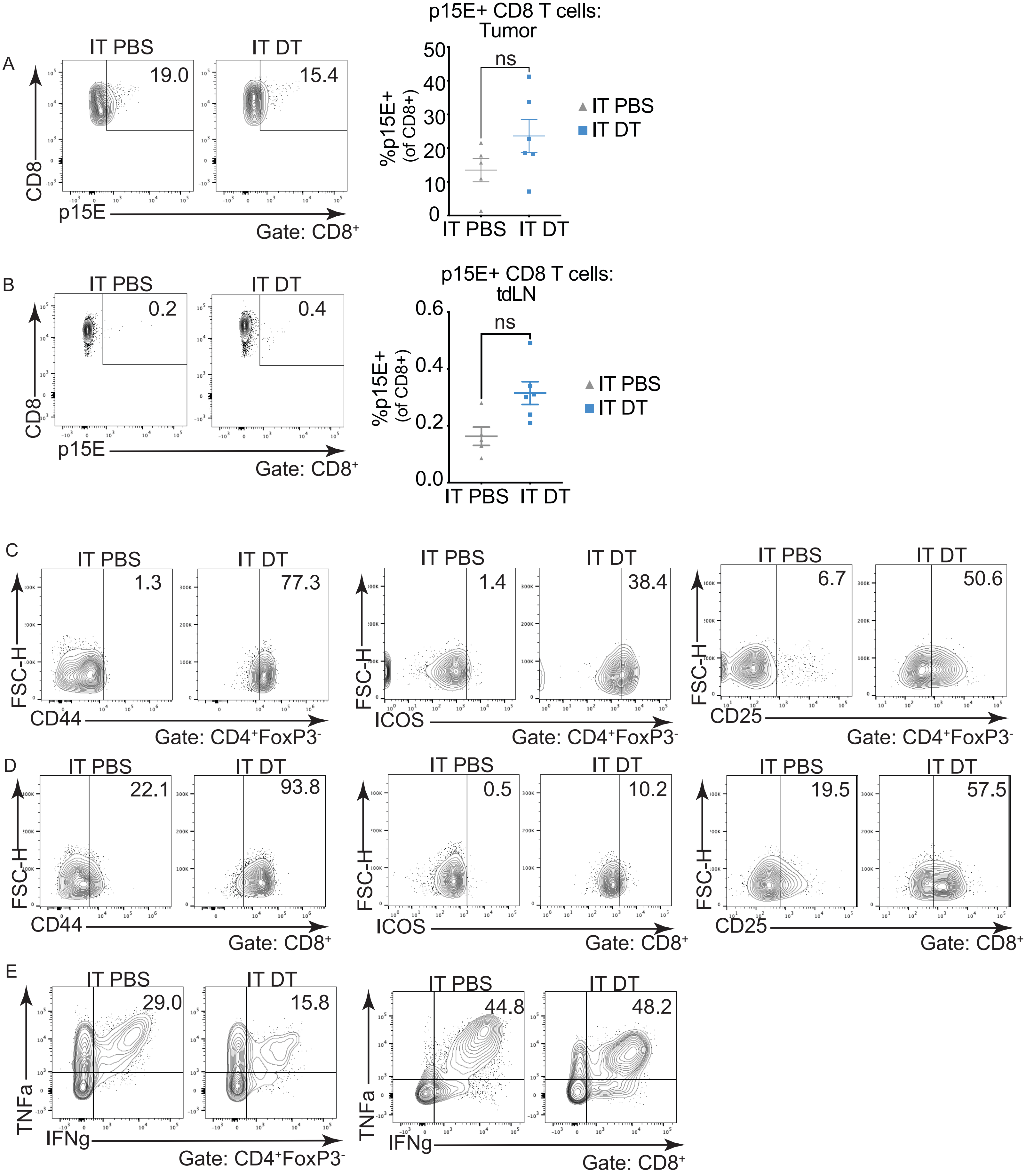

SUPPLEMENTAL FIGURE 3 (cont)

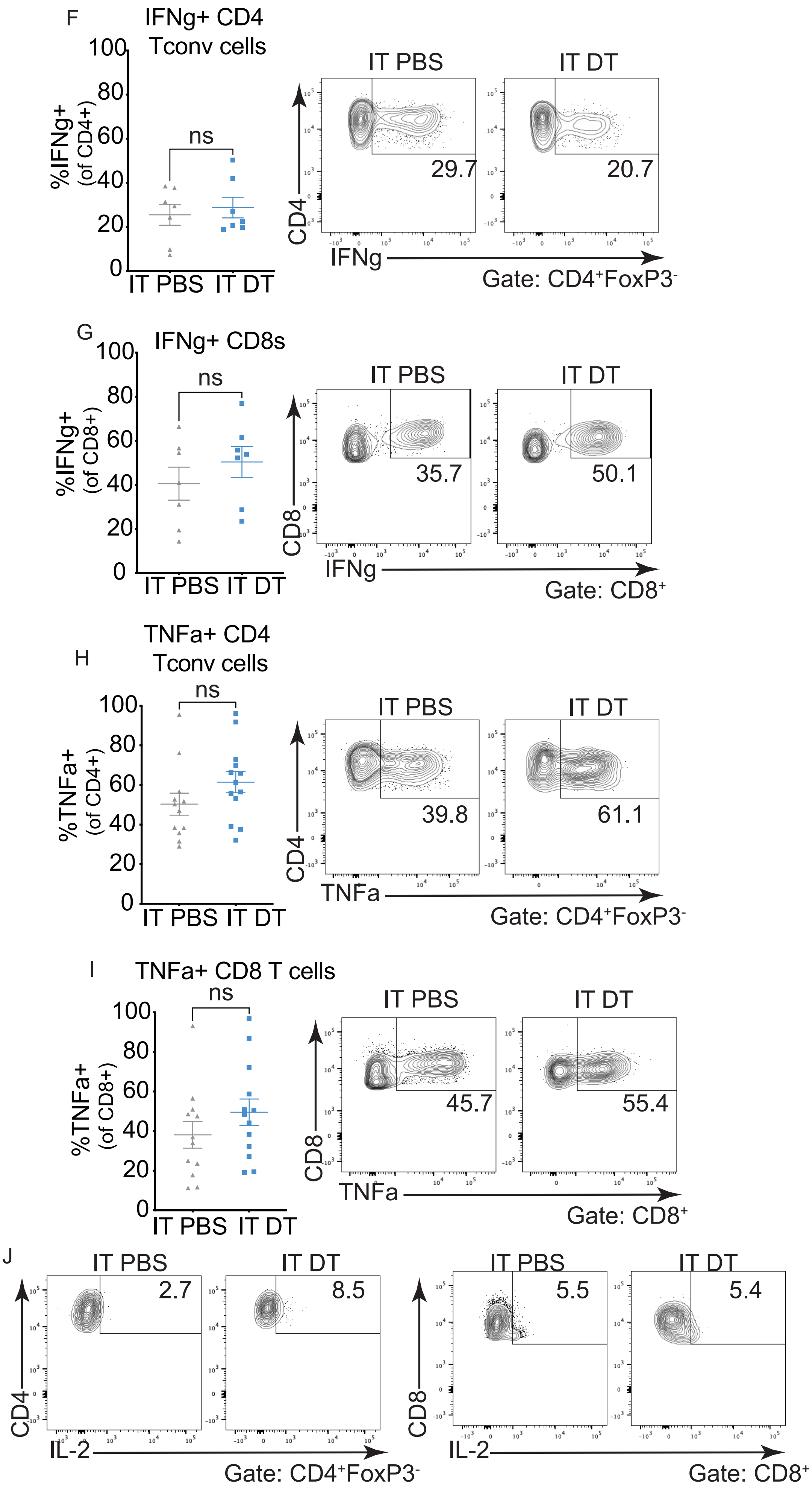

SUPPLEMENTAL FIGURE 3 (cont. II )

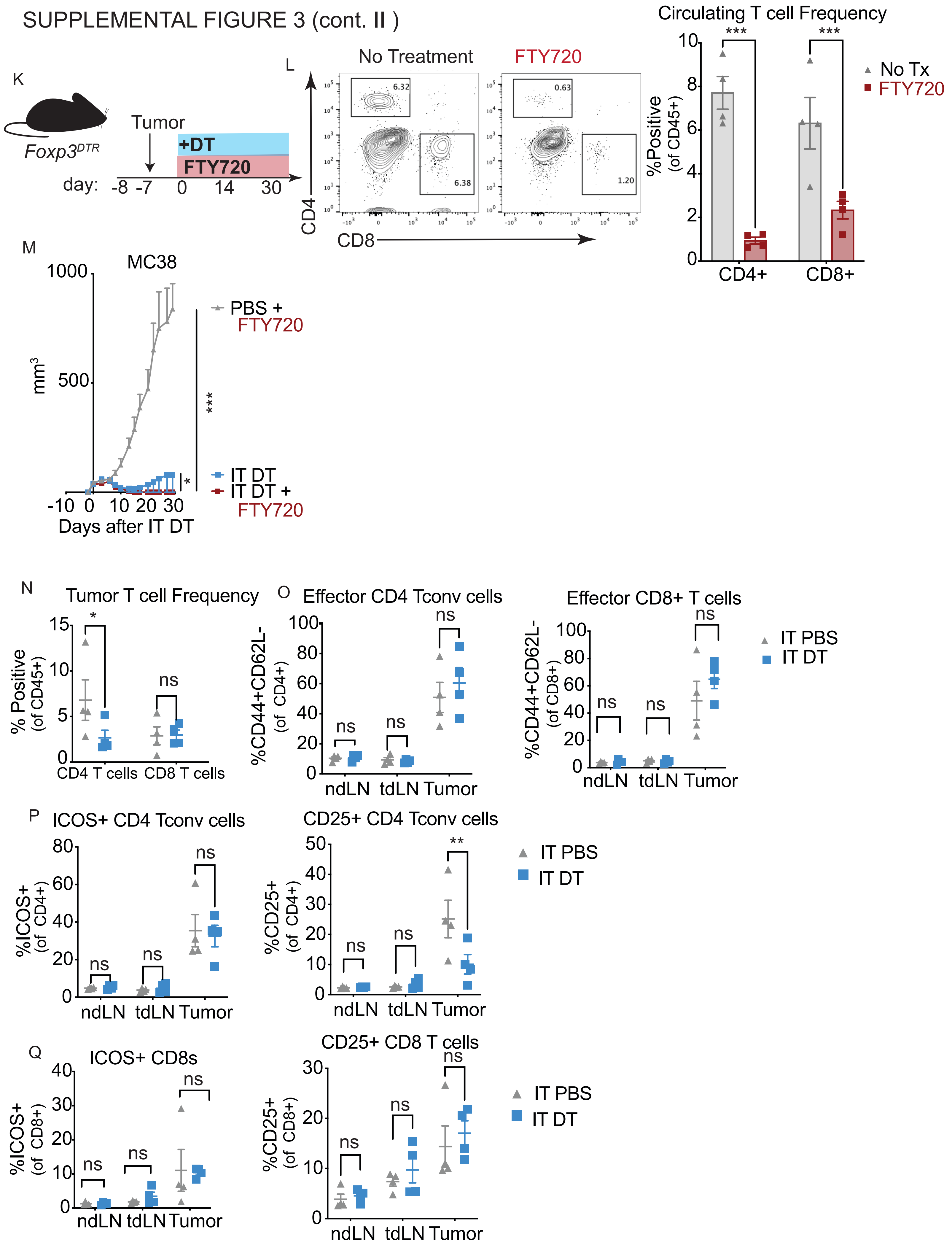

**Supplemental Figure 3. Intratumoral Treg depletion promotes CD8+ and CD4+ T cell activation locally in tumors.**

(A-B) Quantification and representative flow plots of p15E-tetramer+ CD8+ T cells in MC38 tumors or (B) tdLNs of IT PBS or IT DT treated *Foxp3*$^{DTR}$ mice.

(C-D) Representative flow plots of CD44, ICOS or CD25 protein expression in CD4+ Tconv cells or (C) CD8+ T cells.

(E) Coexpression of TNF$\alpha$ and IFN$\gamma$ in CD4+ Tconv cells (left) or CD8+ T cells (right).

(F-G) Quantification and representative flow plots of IFN$\gamma$ expression on CD4+ Tconv cells or (F) CD8+ T cells.

(H) TNF$\alpha$+ expression on CD4+ Tconv cells or (H) CD8+ T cells.

(J) Representative flow plots showing IL-2 production from CD4+ Tconv cells (left) or CD8+ T cells.

(K) Experimental schematic of FTY720 and IT DT dosing.

(L) Flow cytometry quantification of circulating CD4+ and CD8+ T cells in peripheral blood after 14 days of FTY720 treatment.

(M) MC38 tumor growth of FTY720 or IT DT + FTY720 treated mice.

(N) Quantification of tumor T cell frequencies and (O) effector CD4 Tconv or CD8 T cells, 48 hours after IT DT treatment.

(P) ICOS and CD25 expression on CD4+ T cells 48 hours after IT DT treatment.

(Q) ICOS and CD25 expression on CD8+ T cells 48 hours after IT DT treatment.

Data are representative of 2 independent experiments. Data represent means ± SEM; $^*p < 0.05$, $^{**}p < 0.01$ and $^{***}p < 0.001$. A;E-H: Unpaired two-tailed T test. L,M;N-Q: Ordinary two-way ANOVA with Tukey's multiple comparison test. (n=4-7 mice/group).

SUPPLEMENTAL FIGURE 4

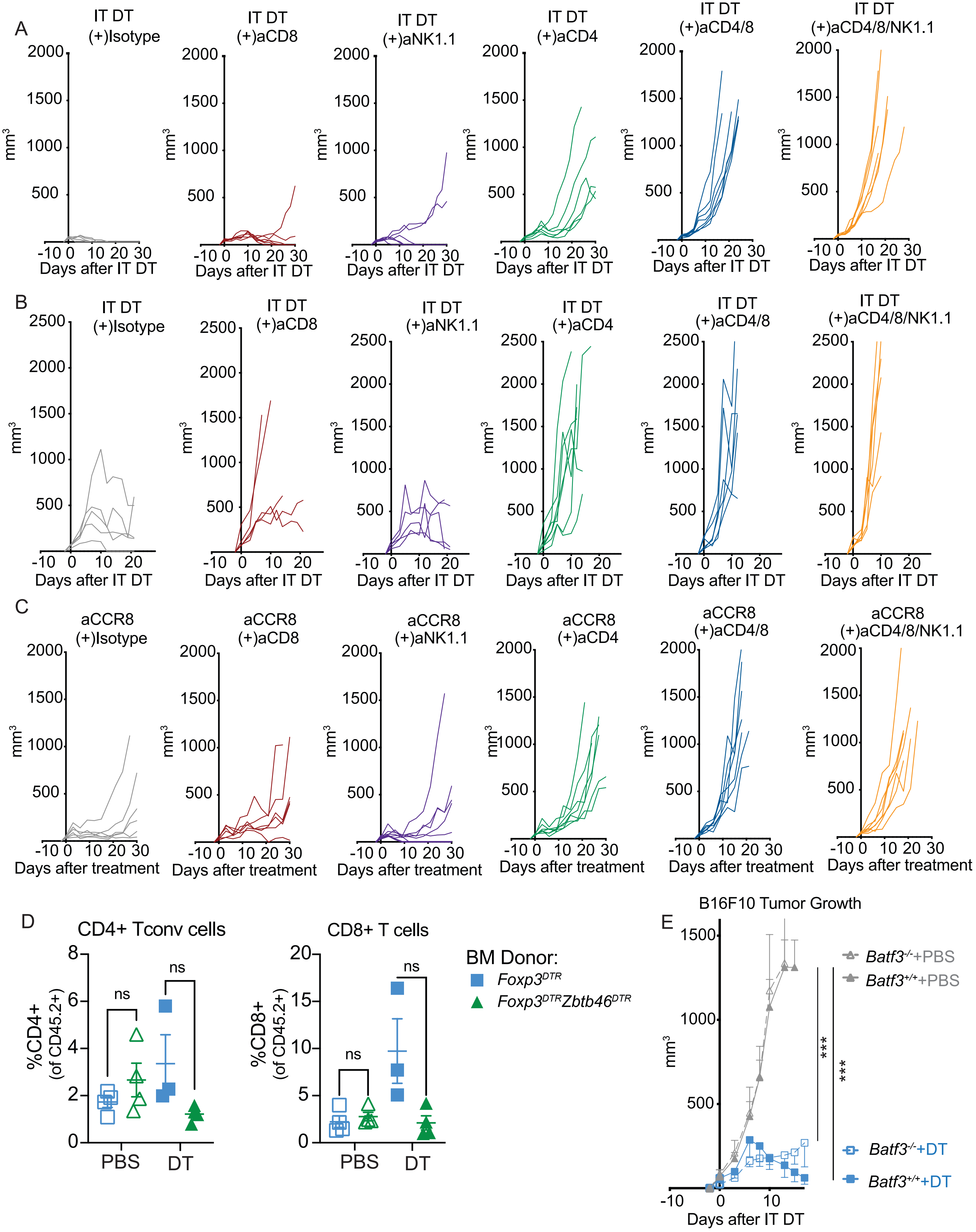

**Supplemental Figure 4. Conventional CD4+ T cells alone  control tumors after IT Treg ablation.**

(A-B) Individual tumor growth curves of MC38 tumors or (B) B16F10 tumors in $Foxp3^{DTR}$ mice after *in vivo* cellular depletion of CD8+ T cells, NK cells, CD4+ T cells or a combination of CD4 & CD8 T cells or CD4, CD8 and NK cells.

(C) Individual tumor growth curves of MC38 tumor bearing mice after *in vivo* administration of anti-CCR8 depleting antibodies and cellular depletion of CD8+ T cells, NK cells, CD4+ T cells or a combination of CD4 & CD8 T cells or CD4, CD8 and NK cells.

(D) Frequencies of conventional CD4+ or CD8+ T cells after in vivo depletion of Tregs and cDCs (green) or Tregs (blue).

(E) B16F10 tumor growth in Treg and cDC1 deficient $Foxp3^{DTR}Batf3^{-/-}$ mice.

Data are representative of 1 independent experiment. Data represent means ± SEM; *$p < 0.05$, **$p < 0.01$ and ***$p < 0.001$. D: Unpaired two-tailed T test. E: Ordinary two-way ANOVA with Tukey's multiple comparison test. (n=4-7 mice/group).

SUPPLEMENTAL FIGURE 5

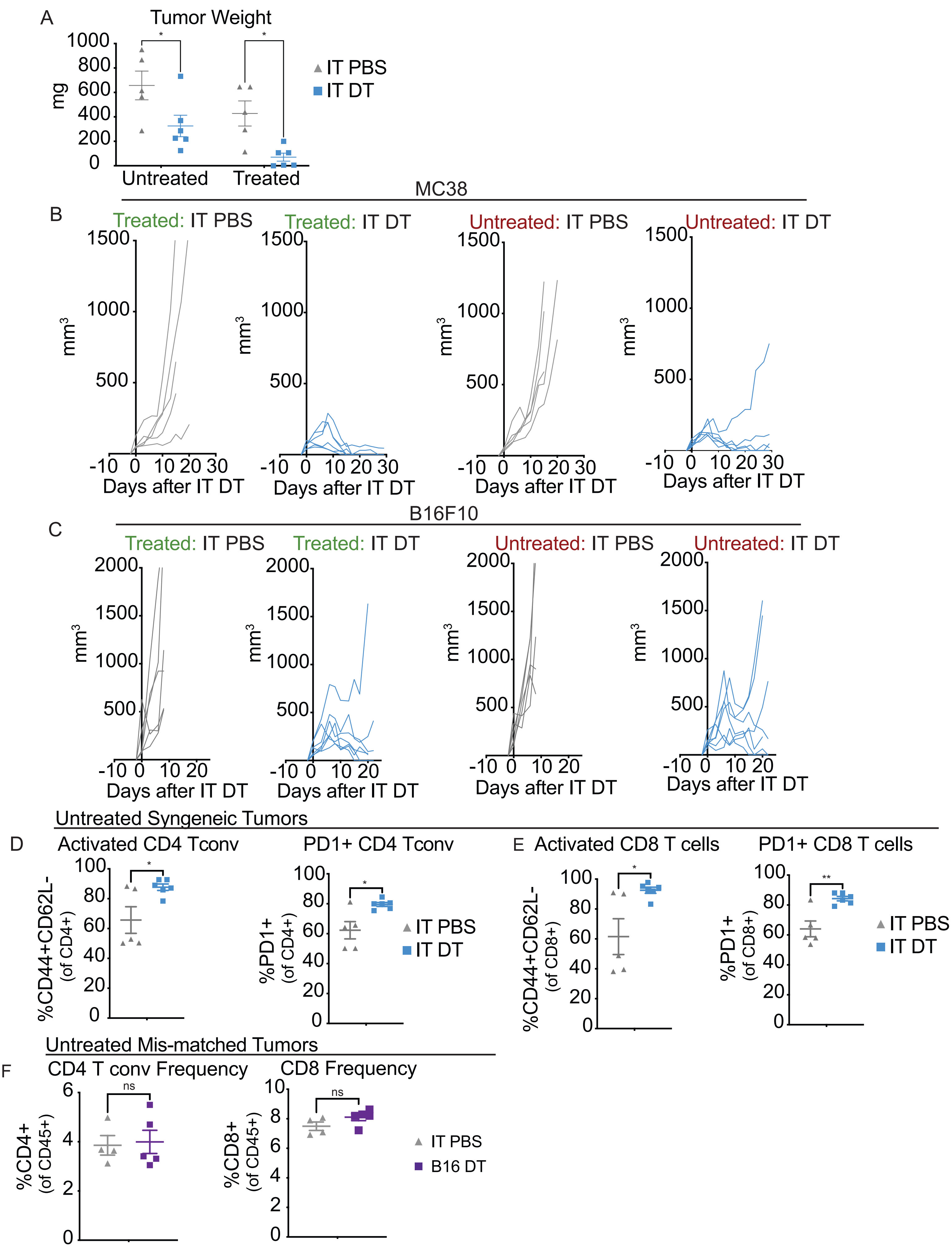

**Supplemental Figure 5. Ablation of IT Tregs in a single tumor unleashes antitumor immunity against distal tumors**

(A) Quantification of MC38 tumor weight of a directly IT DT treated or untreated tumor.

(B-C) Individual tumor growth curves in MC38 dual tumor-bearing *Foxp3*$^{DTR}$ mice or (C)B16F10 tumor-bearing *Foxp3*$^{DTR}$ mice.

(D) Frequencies of activated (CD44+CD62L-) or PD1+ CD8+ T cells in a distal, untreated MC38 tumor.

(E) Frequencies of activated (CD44+CD62L-) or PD1+ CD4+ Tconv cells in a distal, untreated MC38 tumor.

(F) CD4+ Tconv cell frequencies or CD8+ T cell frequencies in a distal, untreated MC38 tumor after IT DT treatment in a mismatched B16F10 tumor.

Data are representative of 2 independent experiments. Data represent means ± SEM; *p < 0.05, **p < 0.01 and ***p < 0.001. A;D-F:Unpaired two-tailed T test. (n=4-7 mice/group).

SUPPLEMENTAL FIGURE 6

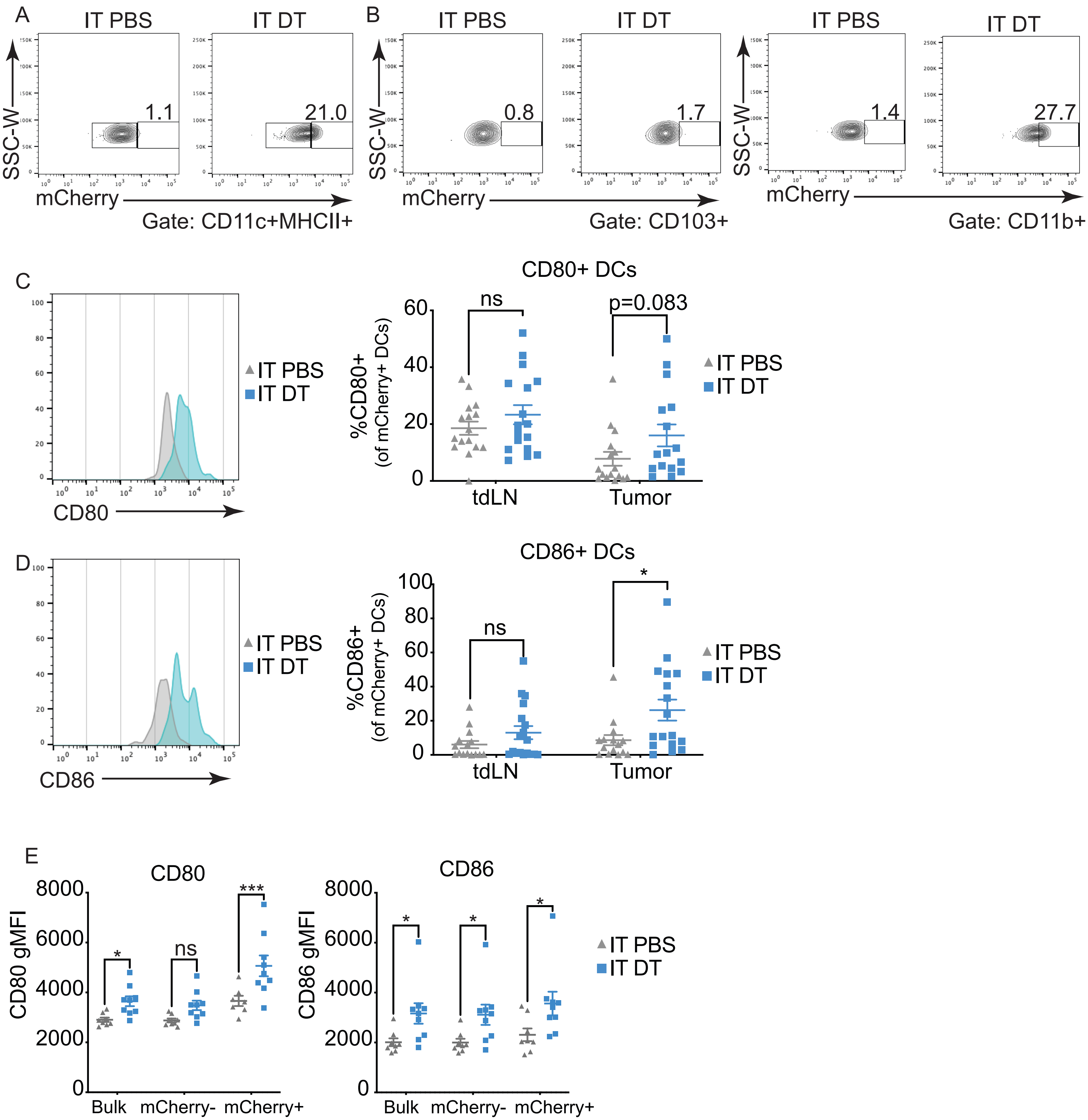

**Supplemental Figure 6. IT Treg ablation enhances tumor antigen acquisition by cDC2s.**
(A) Representative flow plots of mCherry+ tumor antigen+ DCs after IT PBS or IT DT.
(B) Representative flow plots of mCherry+ cDC1 (left) or cDC2 (right) populations after IT PBS or IT DT.
(C-D) Histogram and quantified data demonstrating CD80 or (D) CD86 surface expression on mCherry+ DC populations after IT PBS or IT DT.
(E) Surface expression of CD80 or CD86 on bulk DCs, mCherry- DCs or mCherry+ DCs.
Data are representative of 2 independent experiments. Data represent means ± SEM; *$p < 0.05$, **$p < 0.01$ and ***$p < 0.001$. C-E: Ordinary two-way ANOVA with Tukey's multiple comparison test. (n=9 mice/group).
